# Single-cell transcriptomic analysis unveils spinal motor neuron subtype diversity underpinning the water-to-land transition in vertebrates

**DOI:** 10.1101/2021.09.29.462340

**Authors:** Ee Shan Liau, Suoqin Jin, Yen-Chung Chen, Wei-Szu Liu, Luok Wen Yong, Chang-Tai Tsai, Maëliss Calon, Jr-Kai Yu, Yi-Hsien Su, Stéphane Nedelec, Qing Nie, Jun-An Chen

## Abstract

Spinal motor neurons (MNs) integrate sensory stimuli and brain commands to generate motor movements in vertebrates. Distinct MN populations and their diversity has long been hypothesized to co-evolve with motor circuit to provide the neural basis from undulatory to ambulatory locomotion during aquatic-to-terrestrial transition of vertebrates. However, how these subtypes are evolved remains largely enigmatic. Using single-cell transcriptomics, we investigate heterogeneity in mouse MNs and discover novel segment-specific subtypes. Among limb-innervating MNs, we reveal a diverse neuropeptide code for delineating putative motor pool identities. We further uncovered that axial MNs are subdivided by three conserved and molecularly distinct subpopulations, defined by Satb2, Nr2f2 or Bcl11b expression. Although axial MNs are conserved from cephalochordates to humans, subtype diversity becomes prominent in land animals and appears to continue evolving in humans. Overall, our study provides a unified classification system for spinal MNs and paves the way towards deciphering how neuronal subtypes are evolved.

## Introduction

Motor behaviors are fundamental in animals, enabling basic survival skills such as fight-or-flight to the enormous repertoire of movements underlying complex social interactions. These movements are controlled by spinal cord circuits composed of diverse cardinal neuronal types, which are categorized based on their neurotransmitter types, morphology, location, electrophysiological profiles, connectivity, developmental lineage and molecular profiles^1^. Among these neuronal types, ventral motor neurons (MNs) are the final hub conveying commands from the central nervous system (CNS) to peripherals, accounting for less than 1% of the cells within the spinal cord^2^. Previous studies have reported on MN diversity during development and have provided insights into how these neurons form precise synaptic connections to accommodate the vast heterogeneity of muscular functions^3^.

Within the spinal cord, the cell bodies of MNs innervating specific muscle targets are topographically organized within columnar, divisional, and pool subtypes. The specification of spinal MN subtypes follows a general principle, with morphogens emanating from the dorsoventral [bone morphogenetic proteins (BMP)/Wnt/Sonic Hedgehog (Shh)] and rostrocaudal [retinoic acid (RA)/fibroblast growth factor (FGF)] regions inducing differential sets of transcription factors (TFs) during MN columnar identity consolidation^4^. Specific combinations of columnar subtypes are observed in different body segments and they provide the neural basis for delicate movements. For instance, medial motor column (MMC) MNs spanning all segments control axial muscles and underlie the basic motor patterns observed in both aquatic and land vertebrates. Specifically, axial muscles in aquatic vertebrates facilitate their lateral undulatory swimming patterns, whereas axial muscles in terrestrial vertebrates stabilize the trunk for postural maintenance^5, 6^. Although it has been hypothesized that different subpopulations of MMC neurons may innervate distinct axial muscles^7^, a systematic exploration of axial MN diversity is still lacking and so too is a comparison of MMC neuronal diversity between aquatic and terrestrial vertebrates.

Lateral motor column (LMC) MNs in chick and rodents display even greater complexity than MMC MNs, being divided into ∼60 subpopulations known as motor pools. Each motor pool innervates specific limb muscle groups. Apart from their selective innervating targets, these LMC motor pools are topographically clustered along limb-levels spinal cord segments (brachial and lumbar), and display a unique gene expression profile mainly involving TFs, axon guidance molecules and ligand/receptor genes^3, 8, 9^. It is likely that some motor pool-specific TFs are important in dictating activity of downstream effector genes during the motor pool specification process. For example, combinatorial *Hox* expression within the brachial spinal cord region activates the expression of motor pool-specific markers *Etv4* (*Pea3*), *Runx1* or *Pou3f1* (*Scip*) and subsequent dendritic and axon patterning genes^10^. This combinatorial gene expression thus plays pivotal roles in mediating robust motor pool specification, nerve-muscle connectivity, and sensory-motor circuit formation. Misexpression of these genes was shown to result in altered motor axon arborization and dendritic patterning, inducing impaired motor axon navigation and circuit formation^11–15^. Given that motor pool-specific gene expression drives distinct cell type-specific functions, we were confident that acomprehensive spinal MN transcriptomic profile would illuminate motor pool identity, as well as the intrinsic and extrinsic mechanisms underlying motor pool specification and diversification.

Previous efforts to identify important molecular regulators of motor pools have focused on large muscle groups amenable to retrograde labeling, so only a dozen motor pools have been defined molecularly to date. Although the retrograde labeling technique preserves information on nerve-muscle connectivity, it only allows very minimal starting materials to be collected for sequencing. Moreover, bulk RNA sequencing only reveals averaged expression from a group of MNs, so rare subtypes are likely to be underrepresented or averaged out. To circumvent these issues, single-cell RNA-sequencing (scRNA-seq) was introduced to profile individual cells^16, 17^. This technique enables heterogeneity within cell populations to be characterized in a comprehensive and unbiased manner, with the prospect of generating a cell ‘atlas’ for an entire organism^18, 19^. Single-cell molecular profiling at developmental and adult stages further provide valuable insights into the mechanism of cell type diversification and cell fate decision, as well as the aberrant change of cell populations and gene regulatory network in diseases^20–24^. Moreover, recent state of art development of single-cell multimodal technology integrates gene expression analysis with epitranscriptomes, chromatin accessibilities, proteomes or spatial analysis, which would empower the pursuit for greater understanding to the biology field^25^.

Recent studies have utilized single-cell analysis to demonstrate the diversity of cholinergic neurons in the adult spinal cord^26, 27^, yet the embryonic origin of spinal MN diversity has not been determined. We tackled this topic by using Mnx1-GFP transgenic mice and established a robust optimized method to enrich for and characterize spinal MNs. Clustering analysis based on transcriptome similarity identified known and novel segmental as well as columnar MN subtypes. Among limb MN columns, we reveal 26 distinct MN subtypes corresponding to previously identified and putatively new motor pools. Finally, we provide insights into the unappreciated heterogeneity of axial MNs and explore how these subtypes arose and are delineated in chordates, including humans. Our results may serve as a useful paradigm for investigating multifaceted regulatory mechanisms responsible for neuronal type specification and diversification during development. We further interpret our findings to explain how axial MNs could have evolved during the aquatic-to-terrestrial transition of vertebrates.

## Results

### An enhanced protocol to isolate viable embryonic spinal MNs

The diverse motor behaviors of mammals are regulated by a rich repertoire of MNs distributed along the spinal cord that is generated during embryogenesis (**Fig. 1a**). However, we still lack a complete picture of spinal MN molecular identity in mammalian embryos. To comprehensively resolve spinal MN heterogeneity, we performed scRNA-seq on spinal MNs, either from rostral (brachial and thoracic vertebral segments, from C4 to T3) or caudal (lumbosacral, L1 to S5) regions at embryonic day 13.5 (E13.5) (**Fig. 1b**), a stage when MN subtype diversity is maximized for selective axon targeting and muscle innervation^28^. We enriched for MNs by using an MN-specific fluorescent reporter (Mnx1-GFP) mouse line, and isolated the cell population with the highest GFP intensity (GFP^high^) by means of fluorescence-activated cell sorting (FACS). Our optimized protocol secured ∼7-fold more viable embryonic MNs compared to a traditional trypsin-based primary MN collection protocol (**Extended Data Fig. 1a**, see Methods)^29^. We found that gentle papain-based spinal cord tissue dissociation, followed by FACS using a wide-diameter nozzle (85 μm) and low sheath pressure (45 psi), and then collection of the cell population displaying only the highest GFP intensity achieved an optimal balance between quantity and viability of collected spinal MNs.

**Fig. 1:**
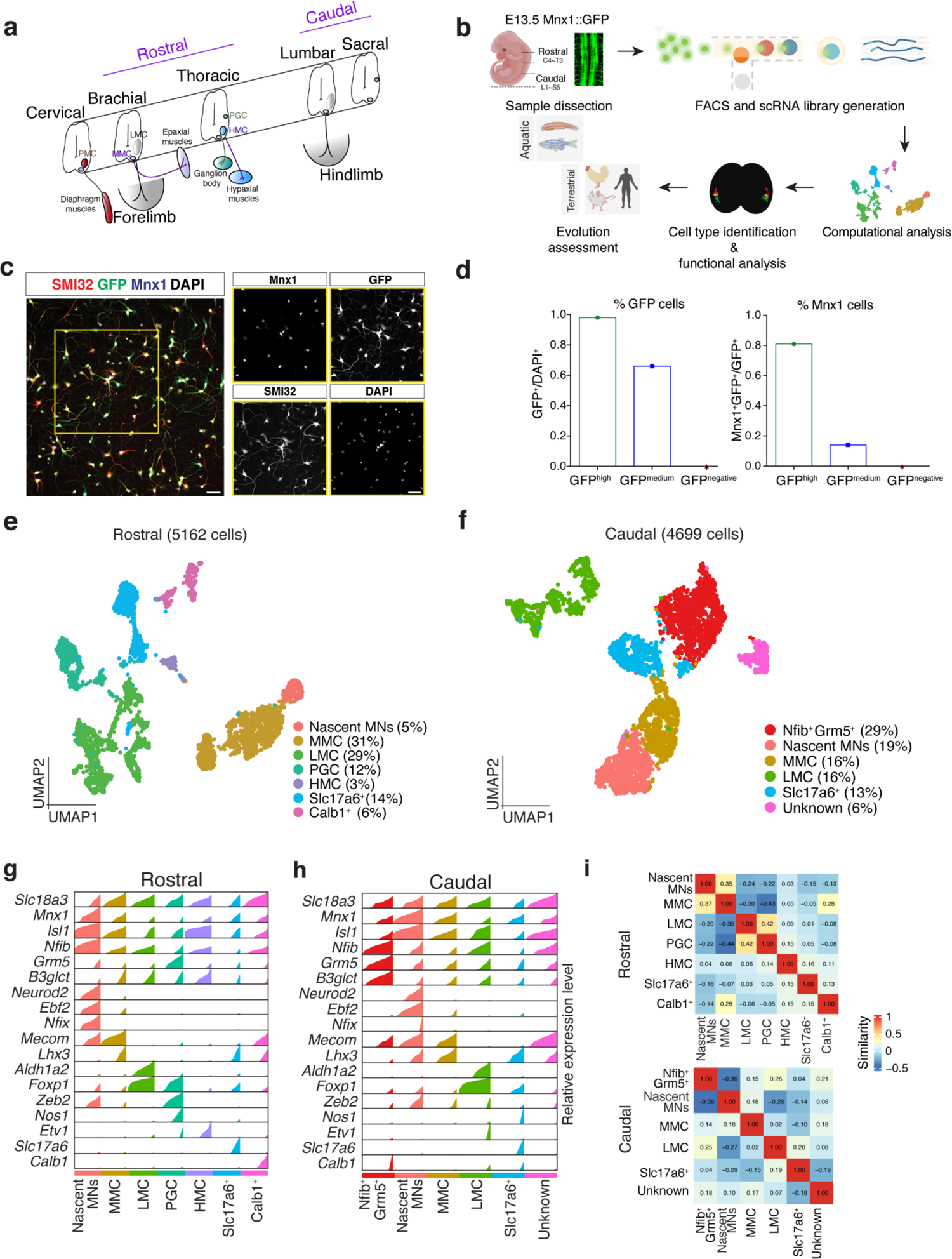
Single-cell transcriptome profiling of E13.5 mouse spinal motor neurons. (a) Illustration of MN subtypes innervating different muscle targets along the rostrocaudal axis of the spinal cord. PMC: phrenic motor column; MMC: medial motor column; LMC: lateral motor column; PGC: preganglionic motor column; HMC: hypaxial motor column. Samples for this study were collected from rostral and caudal segments. (b) Overview of the experimental workflow. (c) Primary MN culture upon dissociation. FACS-collected samples cultured for 48 h were collected for immunostaining (MN markers: SMI32, Mnx1; nuclear marker: DAPI). High-magnification images of individual channels are shown on the right. Scale bar represents 50μm. (d) Quantification of GFP and Mnx1-expressing cells for different samples collected based on GFP intensity (GFP^high^, GFP^medium^ and GFP^negative^) from immunostaining results in (c). (e and f) UMAP plot showing cellular heterogeneity of rostral (e) and caudal (f) samples after removing interneuron and non-neuronal cells. Cell populations are color-coded and cell identities have been annotated based on the expression patterns of known and novel markers. Percentages of each cell population are shown in parentheses. (g and h) High-density bar charts showing relative expression levels of known and newly-identified marker genes (rows) in each cluster (columns) for rostral (g) and caudal (h) MNs, respectively. Within every cluster, cells are divided into ten bins based on increasing expression levels and bar heights correspond to the average expression level of the genes in each bin. (i) Similarity analysis between clusters from rostral (upper) and caudal (bottom) MNs.

Before performing single-cell isolation with a 10x Genomics platform, we performed bulk qPCR and primary culture to verify GFP^high^ cell identity and viability. The GFP^high^ cells displayed *bona fide* MN identity and high expression of hallmark genes for MNs (*Mnx1*, *Isl1*, *Chat*), together with minimal expression of an interneuron marker gene (*En1*) **(Extended Data Fig. 1b**). Expression patterns of rostral (*Hoxc6*^+^) and caudal (*Hoxd10*^+^) markers were in accordance with stringent microdissection and successful separation of spinal segments **(Extended Data Fig. 1c**). Most importantly, sorted GFP^high^ MNs were amenable to culture, exhibited robust neurite outgrowth, and expressed typical MN markers after 2 days (**Fig. 1c, d)**, demonstrating that the spinal MNs we isolated were alive and healthy after our dissection and purification steps. Thus, our customized protocol allows for enrichment of vigorous spinal MNs of known segmental identity for subsequent single-cell transcriptomic profiling.

### Identification of spinal MN subtypes and their novel signatures

We subjected our isolated rostral and caudal Mnx1-GFP^high^ MNs to scRNA-seq. Median numbers of genes detected per cell were 2843 and 2722 for rostral and caudal samples, respectively, and median numbers of unique transcripts were 11,906 and 11,090, respectively (**Extended Data Fig. 1d**). Low-quality cells (e.g., damaged or multiplets) were excluded from analysis by filtering out outliers based on unique molecular identifier (UMI) counts and high percentages of mitochondrial genes **(Extended Data Fig. 2a, d)**. Cells that passed these quality controls were computationally clustered based on transcriptome similarities. Approximately 99% of 10702 rostral or caudal cells were annotated as neurons based on expression of the pan-neuronal markers *Map2* or *Nefl*. Among these neurons, a tiny group of cells (rostral: ∼300 cells; caudal: ∼435 cells) expressed spinal interneuron (IN) markers such as *En1*, *Pou4f1* or *Pax2*, whereas the remaining groups of cells (rostral: 5,162 cells (∼95%); caudal: 4699 cells (∼90%)) expressed cholinergic markers (*Slc18a3* and *Chat*) and spinal MN markers (*Mnx1* or *Isl1*), in agreement with *bona fide* MN identity **(Extended Data Fig. 1d, 2b, c, e, f)**. Unsupervised clustering of MNs using Seurat R package^30^ identified seven and six major clusters among the rostral and caudal sets, respectively **(Fig. 1e and 1f, marker genes in Supplementary Tables 1 and 2)**. We were able to attribute four clusters to known motor columns^3^ by inspecting the expression of established markers: limb-innervating MNs (LMC: *Foxp1*^+^ and *Aldh1a*2^+^), epaxial muscle-innervating MNs (MMC: *Mecom*^+^ and *Lhx3*^+^)^31^, sympathetic ganglion-innervating preganglionic column MNs (PGC: *Nos1*^+^ and *Zeb2*^+^) and hypaxial intercostal muscle-innervating hypaxial motor column MNs (HMC: *Lhx3*^-^, Foxp1^-^, Isl1^+^) **(Fig. 1g and 1h)**. These major columnar MN subtypes largely segregate in a uniform manifold approximation and projection (UMAP) space. Strikingly, five of our clusters could not be assigned to previously documented MN subtypes^32^ (**Fig. 1i** and Extended Data Fig. 3a). We were able to annotate four clusters as nascent MNs (due to their strong expression of neurogenesis-related genes such as *Neurod2* and *Ebf2*)^33^, Mnx1^+^ excitatory interneurons (*Slc17a6*^+^)^34^, and two new segmentally-restricted subtypes, i.e., the Calb1^+^ cluster from the rostral segment that expresses *Mnx1* and *Calb1*, and the Nfib^+^Grm5^+^ cluster from the caudal segment (**Fig. 1g, h)**.

**Fig. 2:**
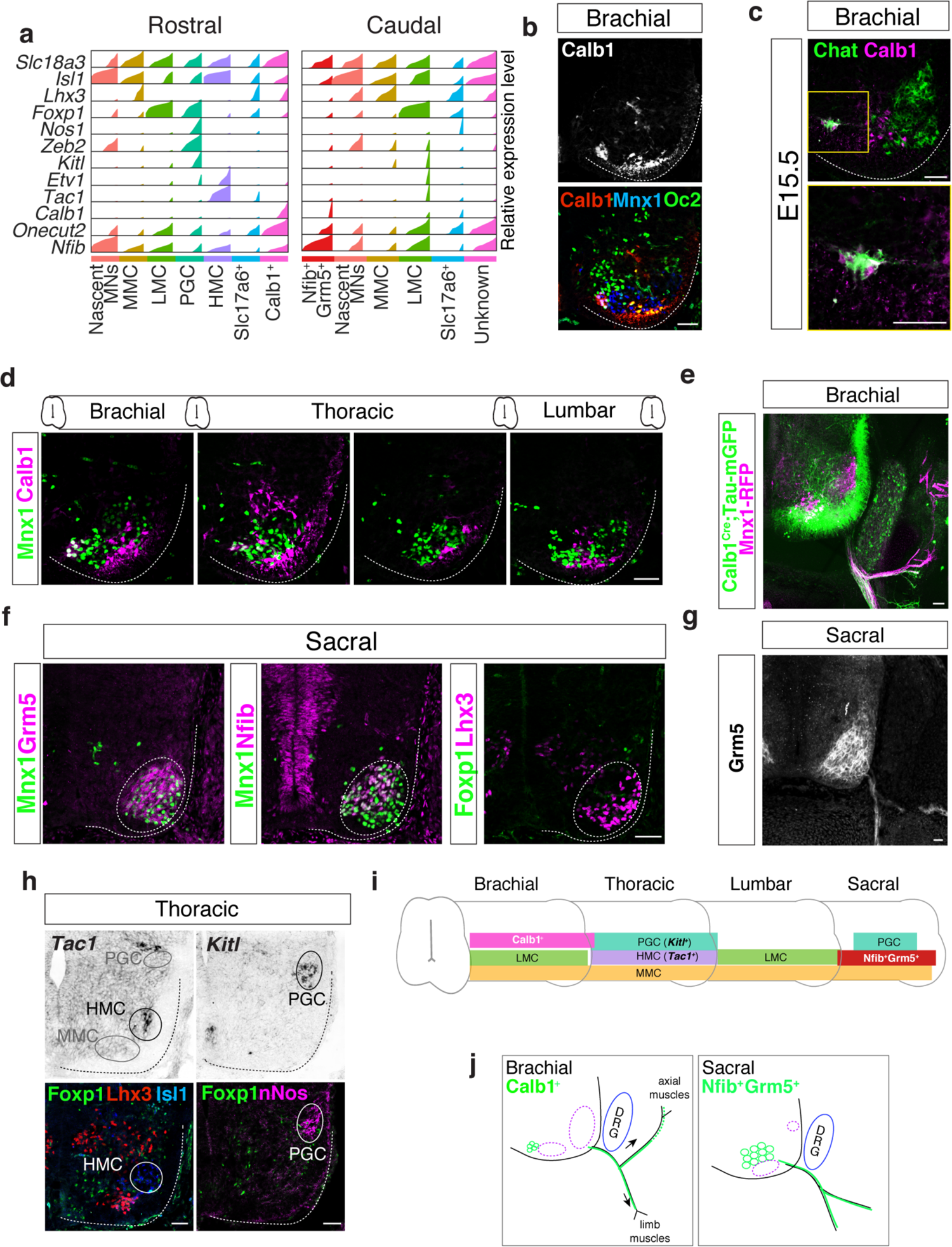
Identification of previously unreported MN subtypes and novel markers. (a) High-density bar charts showing relative expression levels of markers enriched in the PGC (*Kitl*), HMC (*Tac1*), Calb1^+^ (*Calb1* and *Onecut2*) and Nfib^+^Grm5^+^ (*Nfib*) clusters. (b) Immunostaining reveals ventromedial positioning of the *Calb1^+^Onecut2^+^Mnx1^+^* subpopulation in the E13.5 mouse spinal cord. (c) At E15.5, immunostaining shows Chat expression in Calb1^+^ subpopulation. A high-magnification image (yellow box) is also shown. (d) *Calb1^+^Mnx1^+^* cells exhibit a gradient of rostrocaudal expression. (e) Calb1^Cre^ mice were crossed with a Tau-mGFP; Mnx1-RFP double reporter to indelibly label axonal trajectories of Calb1^+^ neurons. Overlapping GFP/RFP pixels indicate that the axonal bundles of Calb1^+^ neurons project towards the peripherals. (f) Immunostaining of Mnx1 with Nfib or Grm5 and Foxp1 with Lhx3 in adjacent sacral spinal cord sections (E13.5). Foxp1 and Lhx3 label sacral PGC and MMC MNs, respectively. Dashed circles define the Mnx1^+^ regions. (g) Immunostaining reveals Grm5 in axons projecting to peripherals. (h) *In situ* hybridization of novel markers *Tac1* and *Kitl* identified in HMC and PGC MNs, respectively. Immunostaining for known subtype markers (HMC: Isl1^+^ Foxp1^-^ Lhx3^-^; PGC: Foxp1^+^ nNos^+^) was performed on adjacent slides. (i) Schematic illustration of MN subtypes identified previously and in the current study. (j) Schematic depictions of the Calb1^+^ and Nfib^+^Grm5^+^ clusters with their projecting axons. Green circles and lines represent cell bodies and axon projections of novel clusters identified in this study. Purple circles mark known MN subtypes. DRG = dorsal root ganglion. In b-d, f, h, white dashed lines outline the spinal cord boundary and dashed circles define MN positions. MN subtypes are indicated in h. Scale bars of all images represent 50 μm.

**Fig. 3:**
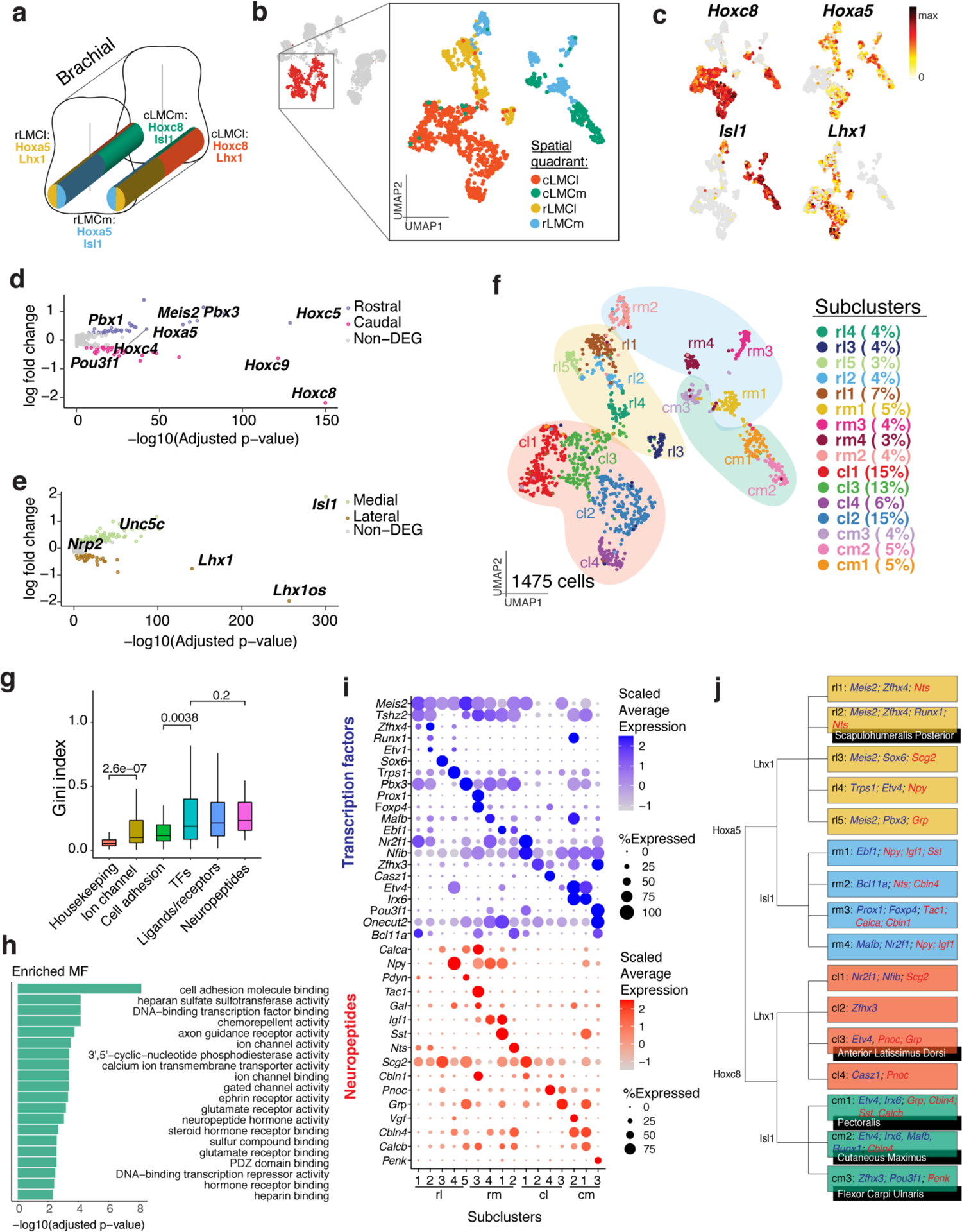
Delineation of molecular diversity in limb MNs by differential expression of transcription factors and neuropeptides. (a) Schematic of limb LMC MN subtypes based on their cell body positions in the ventral horn of the brachial spinal cord. Cells residing in the brachial segment are divided into rostral, caudal, lateral and medial regions and are represented by expression of *Hoxa5*, *Hoxc8*, *Lhx1* and *Isl1*, respectively. (b) UMAP plot of cells from brachial LMC cluster in Fig. 1E (red). LMC MNs were grouped into four “spatial quadrants”, including rLMCl (rostral lateral, yellow), rLMCm (rostral medial, blue), cLMCl (caudal lateral, red) and cLMCm (caudal medial, green), based on the expression patterns of *Hox* genes and LIM homeodomain factors. (c) Expression pattern of rostral (*Hoxa5*), caudal (*Hoxc8*), medial (*Isl1*), and lateral (*Lhx1*) markers on a UMAP plot, reflecting the corresponding “spatial quadrant” identities in (b). (d and e) Volcano plots showing the differentially expressed genes (DEGs) between rostral and caudal (d), and medial and lateral (e) brachial LMC neurons. Genes with an adjusted p-value <0.05 and log fold-change >0.25 were deemed differentially expressed and are color-labeled. Other non-significant genes are indicated by grey dots. DEGs of particular interest are labeled. (a) (f) UMAP plot of brachial LMC subclusters. Cell proportions for each subcluster are indicated as percentages. The four “spatial quadrants” are shaded to match the colors in (b). (b) (g) Comparison of the gene expression variations grouped based on their functional terms within LMC MNs, as accessed by the Gini index. Housekeeping genes were used as a reference. P-values are from two-tailed Wilcoxon rank sum test. (c) (h) Gene ontology enrichment of molecular functions of DEGs across brachial LMC neurons. Terms of interest in this study are highlighted in bold. (i) Dot-plot showing expression patterns of representative markers (rows) from the top differentially-expressed TFs and neuropeptides among LMC subclusters (columns). The size of each circle reflects the percentage of cells expressing the genes, and color intensity reflects scaled average expression for each gene. (d) (j) Summary of TF (blue font) and neuropeptide (red font) combinatorial codes that define the four LMC “spatial quadrant” clusters (boxes shaded as in Fig. 3b) and the sixteen subclusters. Known motor pools are annotated with their corresponding muscle targets in the black boxes.

To further characterize these two novel cell populations, i.e., the rostral Calb1^+^ and caudal Nfib^+^Grm5^+^ clusters, we performed immunostaining for marker proteins on E13.5 spinal cord sections (**Fig. 2a, b)**. The rostral Calb1^+^ cluster was so named according to its expression of *Calb1*, which encodes a calcium-binding protein and is reported to be a distinguishing feature of Renshaw cells, an early-born subtype of V1 interneurons^35^. Cells co-expressing both Mnx1 and Calb1 were found at the most ventromedial region of the spinal cord, distinct from Calb1^+^-only Renshaw cells positioned ventrolateral to MNs (**Fig. 2b**). This cluster of *Mnx1^+^Calb1^+^* cells displayed cholinergic neurotransmitter identity, as revealed by their co-expression of the *Chat*, *Slc5a7* and *Slc18a3* marker genes in our scRNA-seq dataset. Immunostaining at E15.5 demonstrated that Calb1^+^ MNs co-expressed Chat (**Fig. 2c**). The *Mnx1^+^Chat^+^Calb1^+^* cluster was only observed in the brachial and upper thoracic segments, implying potential muscle group innervation specifically in the upper body (**Fig. 2d**). To examine if, like other MN subtypes, the *Mnx1^+^Chat^+^Calb1^+^* neuron population extends its axons outside the spinal cord, we monitored axon trajectories from Calb1^Cre^;Tau-mGFP together with Mnx1-RFP reporter embryos. Strikingly, we observed that the axons of *Mnx1^+^Chat^+^Calb1^+^* neurons extended beyond the spinal cord (**Fig. 2e**). Therefore, we have identified a novel brachial MN column manifesting *Mnx1^+^Chat^+^Calb1^+^* expression that extends its axons out from the spinal cord.

In contrast, the Nfib^+^Grm5^+^ cluster was identified exclusively in the caudal sample (**Fig. 1f and Extended Data Fig. 3c**). Nfib is dynamically expressed in many MN subtypes, so we used an additional marker, *Grm5*, to verify their sacral distribution in embryonic spinal cord **(Extended Data Fig. 3b)**. Immunostaining indeed validated that Nfib^+^Grm5*^+^* co-expressing neurons were only observed in sacral spinal segments (Foxp1^-^ Hoxd10^-^) (**Fig. 2h**; Extended Data Fig. 3c). The Nfib^+^Grm5^+^ population was distinct from Foxp1^+^ and Lhx3^+^ cells, indicating that they are not sacral PGC or MMC MNs (**Fig. 2f**). Moreover, the Nfib^+^Grm5^+^ population appeared to project their axons ventrally to peripherals (**Fig. 2g**).

In addition to our novel MN subtypes, we compared differentially expressed genes (DEGs) among known columnar MN subtypes to reveal novel molecular signatures. HMC MNs were thought to be lack a salient expression hallmark^36^, yet we identified a series of marker genes, including *Tac1* (which encodes the tachykinin peptide hormone family, neurokinin A, substance P, neuropeptide K and neuropeptide gamma) (**Fig. 2a**). *Tac1* was previously described as being expressed by pain-related projection neurons and interneurons in the spinal cord^37, 38^, but to our knowledge its expression has not been reported before for MNs. *In situ* hybridization validated selective expression of *Tac1* in thoracic HMC MNs, representing a novel HMC genetic marker (**Fig. 2h, left)**. Similarly, we have identified *Kitl* as a novel marker of PGC MNs and verified our findings by means of *in situ* hybridization (**Fig. 2h, right)**. *Kitl* encodes ligand for the versatile Kit pathway involved in axon outgrowth, cell migration and cell survival^39, 40^.

Next, we compared LMC and MMC neurons, the two primary MN subtypes that project differentially to limb and axial muscles, respectively. Apart from using TFs (LMC: *Foxp1* and MMC: *Mecom/Lhx4*) and cell surface receptors (MMC: *Lifr/Il6st*) to distinguish these subtypes, which have been reported to regulate cell fate specification and survival^41^, we also uncovered selective expression of IgLON neural adhesion molecules within these neurons **(Extended Data Fig. 4, Supplementary Tables 3 and 4**). For instance, *Ntm* expression discriminates LMC from *Lsamp-*expressing MMC neurons in E13.5 brachial spinal cord **(Extended Data Fig. 4b, c)**. IgLON neural adhesion molecules serve to regulate axon pathfinding, dendritic arborization and synaptogenesis^42–44^, and selective expression of respective genes might contribute to the differences in neuronal pathfinding and circuit wiring between limb-innervating LMC and axial muscle-innervating MMC MNs. In addition, by identifying these markers not only do we provide candidates that could tune neural wiring and axon pathfinding, but they also represent novel targets for affinity-based subtype isolation, circumventing the need for FACS that can be stressful to cells.

**Fig. 4:**
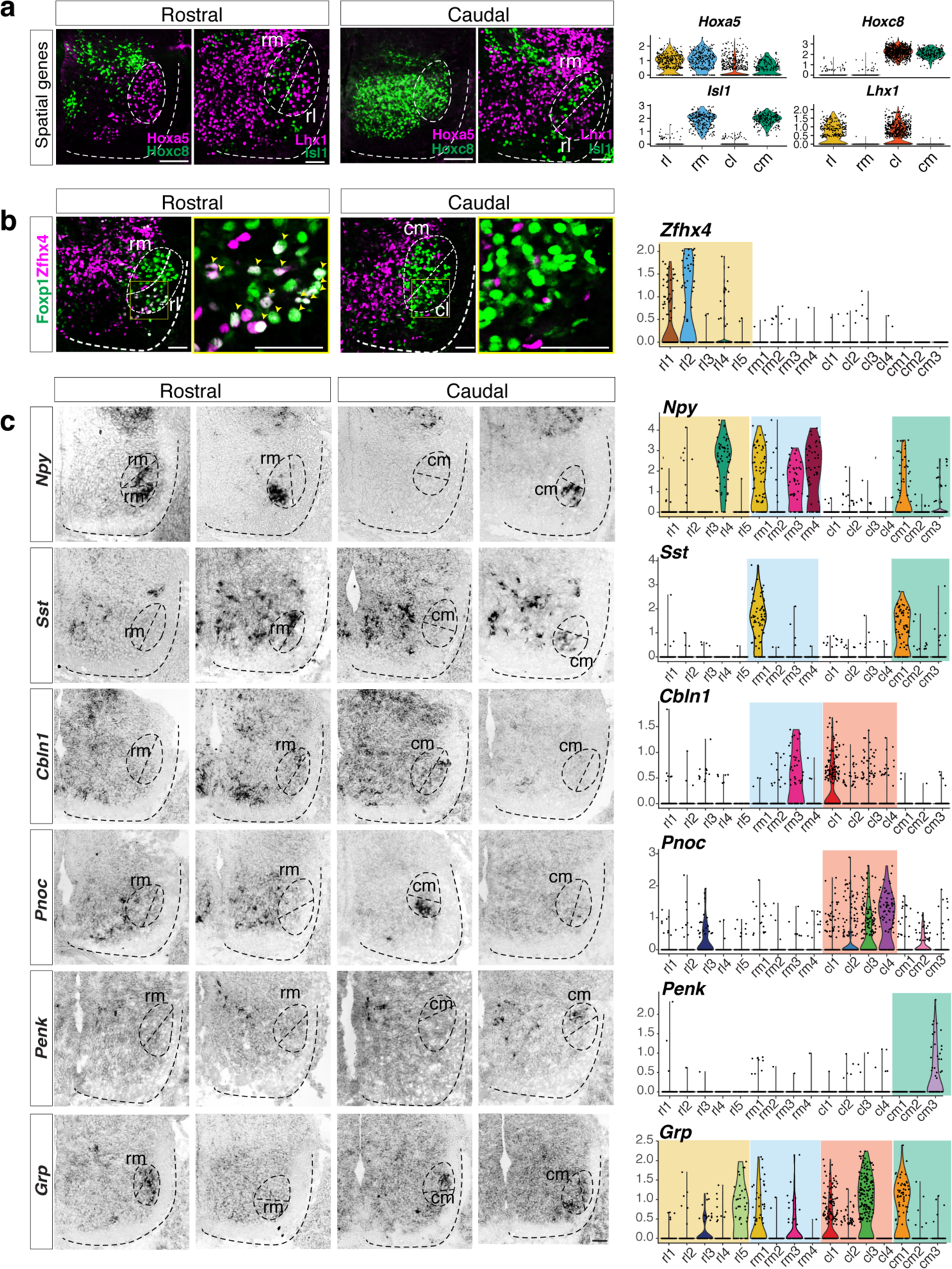
Validation of cluster-specific TFs and neuropeptides defining LMC MN diversity. (a) Representative images of immunostaining for spatial genes on adjacent slides to the marker gene staining shown in panels (b) and (c). *Hoxa5*, *Hoxc8*, *Isl1* and *Lhx1* expression distinguish rostral (r), caudal (c), medial (m) and lateral (l) LMC MNs, respectively. Violin plots (right) show expression of these genes in the brachial LMC “spatial quadrants” established from our scRNA-seq data. (b) Immunostaining (left) and scRNA expression (right) of the transcription factor *Zfhx4* in rostral lateral LMC MNs (rl1 and rl2). Yellow arrowheads in the high-magnification image of the lateral region of LMC MNs indicate expression of Zfhx4. Violin plot (right) shows expression of *Zfhx4* across 16 subclusters of LMC MNs. (c) *In situ* hybridization (ISH) (left) and scRNA expression (violin plots, right) of neuropeptides in LMC MNs. The spatial information demarcated by the dashed lines in (b) and (c) was annotated based on the immunostaining for *Hox* genes (*Hoxa5*/*Hoxc8*) and LIM homeodomain TFs (Isl1/Lhx1) on adjacent slides, as shown in (a). Scale bar depicts 50 μm. r = rostral; c = caudal; m = medial; l = lateral.

Collectively, our scRNA-seq data have not only identified all previously known spinal MN columnar subtypes with novel genetic markers, but also uncovered previously undescribed MN subtypes (**Fig. 2i, j)**.

### Cellular heterogeneity and differential expression of neuropeptides within LMC MNs

LMC MNs are specific to brachial and lumbar levels in the spinal cord. Given that within brachial LMC MNs, previous studies have demonstrated that Hoxa5/Hoxc8 proteins define rostrocaudal (rc) brachial LMC MNs identity^10^, while Isl1/Lhx1 distinguish LMC MNs in mediolateral (ml) positions^45^ (**Fig. 3a**), we analyzed these four cardinal TFs for the 1,475 brachial LMC MNs to assign them in three-dimensional ‘spatial quadrants’ **(Fig. 3b and 3c**). We then examined DEGs between the cells from different ‘spatial quadrants’. We identified a number of Hox cofactors (*Pbx1*, *Pbx3*, *Meis1* and *Meis2*) that were differentially expressed between the rostral (Hoxa5^+^, Hoxc8^-^) and caudal (Hoxc8^+^) brachial clusters^46^, (**Fig. 3d, Supplementary Table 5**). To compare lateral and medial clusters, we observed that *Nrp2* and *Unc5c* were differentially expressed in the medial clusters, consistent with their distributions shown in previous studies^15, 47^ (**Fig. 3e, Supplementary Table 6**). These results corroborate LMC MN assignment to spatial quadrants, faithfully recapitulating MN distributions *in vivo*.

Next, we clustered cells into 16 subclusters distributed across the four spatial quadrants (namely <u>rm</u>: rostral medial, <u>rl</u>: rostral lateral, <u>cm</u>: caudal medial, cl: caudal lateral) (**Fig. 3f**). To prevent over-clustering and determine a suitable number of subclusters, we proposed a multi-resolution ensemble strategy based on spectral graph theory and ensured that we captured distinct clusters at the transcriptomic level (**Extended Data Fig. 5a**; see Methods). An analysis of DEGs demonstrated that each subcluster was represented by a distinct combination of marker genes **(Extended Data Fig. 5b, c, Supplementary Table 7**).

**Fig. 5:**
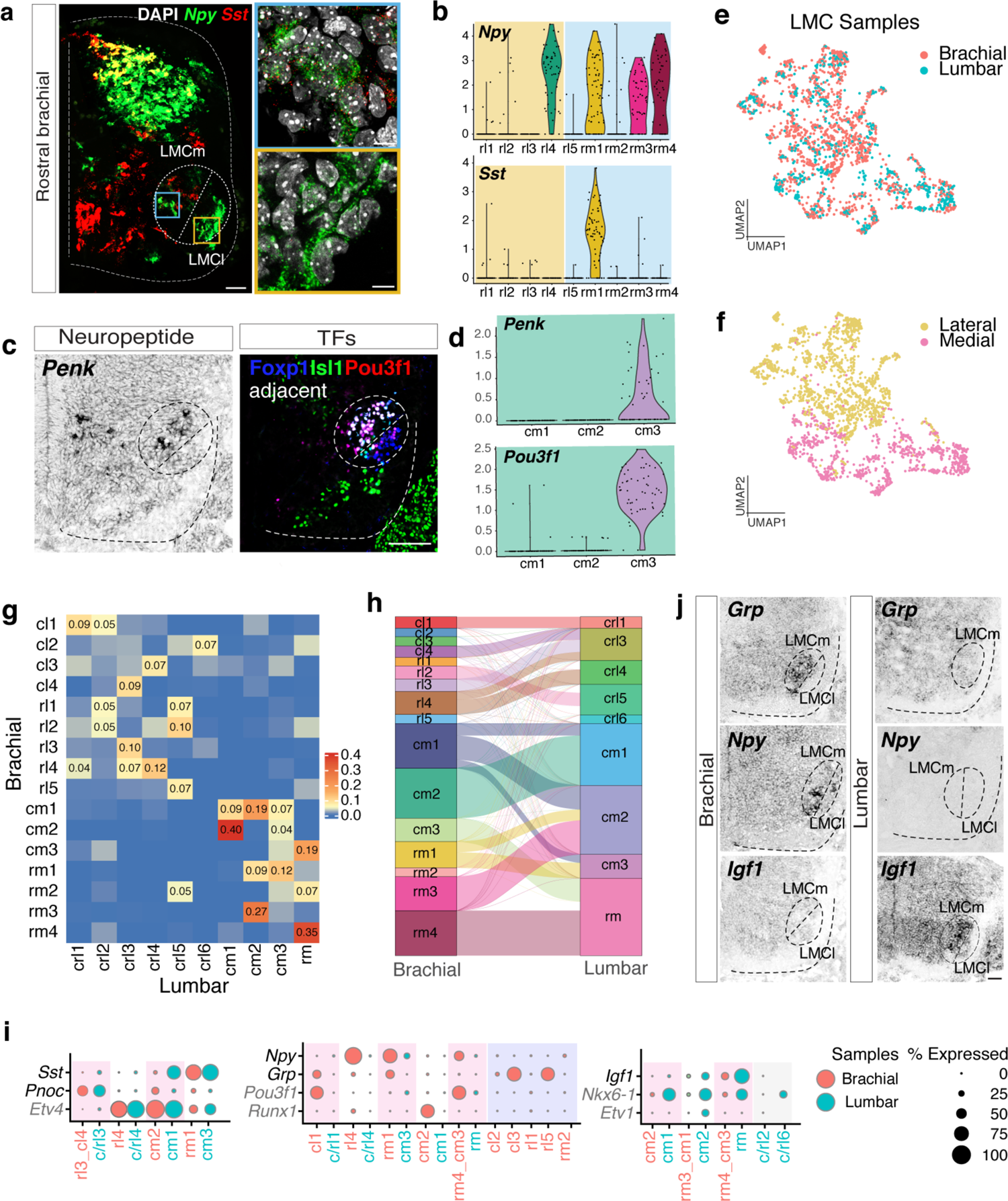
Divergent LMC diversity in brachial and lumbar segments. (a) RNAscope-based fluorescence *in situ* hybridization of *Sst* and *Npy* reveals co-expression in the rm1 subcluster, whereas only *Npy* is expressed in the rl4 cluster. High-magnification images of the rm1 (blue box) and rl4 (yellow box) subclusters are shown on the right. Scale bars for the spinal cord hemi-section and high-magnification images represent 50 μm and 10μm, respectively. (b) Violin plot showing normalized expression of *Npy* and *Sst* in rostral LMC subclusters. (c and d) Representative image of LMC subclusters displaying combinatorial expression of a TF (*Pou3f1*) and a neuropeptide (*Penk*) in caudal medial LMC neurons, supporting the single-cell expression pattern shown in (d). Adjacent slides were used. Scale bar represents 50 μm. (a) (e) Joint UMAP visualization of rostral and caudal LMC MNs showing that cells from both samples could be integrated without confounding technical artifacts. (b) (f) LMC MNs with medial and lateral identities segregated on the joint UMAP space. (c) (g) Similarity analysis between LMC subclusters from rostral and caudal segments, as reflected by the heatmap, which was quantified by the proportion of overlapping mutual nearest neighbors. (d) (h) Alluvial diagram intuitively illustrate the relatedness between brachial and lumbar LMC subclusters. Line thickness reflects level of similarity between subclusters, and the size of each box reflects total similarity. Lines with similarities < 0.09 have been omitted. Some, but not all, subclusters exhibit high similarity between the rostral and caudal segments, indicating that certain subclusters show segmental specificity. (i) Combined dot-plot showing marker gene expression (color-coded along Y axis: grey for known motor pool markers; black for neuropeptides) in LMC subclusters (X axis) from a merged dataset. Three categories are highlighted: (left) selected genes expressed in both brachial and lumbar subclusters; (middle) genes preferentially expressed in brachial segments; (right) genes preferentially expressed in lumbar segments. LMC subclusters in different segments have been grouped based on similarity data from (g) and (h). Circle size indicates the percentage of cells expressing the genes. (e) (j) ISH demonstrating that *Grp* and *Npy* are mainly expressed in brachial segments, whereas expression of *Igf1* is more enriched in lumbar LMC MNs, supporting the single-cell gene expression data presented in (i). Scale bar represents 50 μm.

Consistent with previous studies^48–50^, TFs, cell adhesion molecules, axon guidance molecules, receptors and ion channels topped the list of the most variable genes in brachial LMC MNs for functions relating to cell fate specification, motor pool sorting and positioning, axon pathfinding and synaptogenesis **(Fig. 3g, 3h and Extended Data Fig. 5d, Supplementary Table 8**). Strikingly, neuropeptides also stood out as a prominent gene term, indicative of plausible cluster-specific expression.

Thereafter, we sought to identify each subcluster from our scRNA-seq dataset based on their respective marker genes. We observed expression of known motor pool markers that corresponded to MNs innervating a single specific muscle group. For instance, *Etv4* (*Pea3*), *Runx1* and *Pou3f1* (*Scip*) were specific to the cm1/2 (caudal medial 1/2), rl2 (rostral lateral 2) and cm3 (caudal medial 3) subclusters, respectively **(Fig. 3i and 3j**). Accordingly, we postulated that some of these LMC subclusters may reflect specific motor pools, so they could be annotated with their corresponding innervating muscle groups^10, 51^. For example, the cm1 subcluster expressed *Etv4*, *Sema3e* and *Cdh8*, but not *Cdh7* **(Extended Data Fig. 5e**), thus representing the cutaneous maximus (CM) motor pool^9^. In contrast, another *Etv4*-expressing cluster (cm2) lacking *Runx1* expression distinguishes the pectoralis-innervating motor pool^52^, whereas the *Pou3f1*-expressing cm3 subcluster represents the median and ulnar nerves innervating the flexor carpi ulnaris (FCU) muscle^10, 53^.

Remarkably, the remaining subclusters have not been documented previously, so we attempted to identify their molecular identities. To do so, we focused on genes encoding TFs, cell adhesion molecules, ligands and receptors, which are hallmark genes reported to contribute to desired molecular and functional diversification among neurons^50, 54^. Using a TF as an example, we identified enriched *Zfhx4* expression in a subset of limb MNs (Zfhx4: rl1 and rl2) (**Fig. 4a, b)**. Based on our *in vivo* validations, Zfhx4 expression is restricted to a subpopulation of LMC MNs in the rostral lateral region of the brachial zone, resembling Zfh2 (a Drosophila ortholog of Zfhx4) patterning that specifies late ventral nerve cord neurons^55^.

We observed selective expression of *Zfhx4* in specific LMC MNs, indicating a plausible yet unappreciated role in delineating motor pools. Among the receptor-ligand subfamily, we noted a cohort of neuropeptides are differentially expressed between LMC MN subclusters (**Fig. 3h, i)**. Neuropeptides are small proteins produced by neurons that usually act on G protein-coupled receptors, which can function via autocrine and/or paracrine signaling with neurotransmitters in a single neuron type to modulate and expand neuronal function^56^. Based on single-cell transcriptomic studies of the CNS, neuropeptides can be used to group neuronal types in the brain and dorsal horn of the spinal cord^57, 58^. We found that a host of neuropeptides, including *Npy, Sst, Grp, Cbln1, Cbln4, Pnoc* and *Penk*, were highly enriched and spatially partitioned in different subsets of LMC MNs (**Fig. 4c**). We also noted that distinct LMC MN subtypes expressed specific combinations of neuropeptides. Our analysis predicted that *Sst* and *Npy* co-expression could distinguish the rm1 cluster from the *Npy*-only rl4 cluster, which was validated *in vivo* using RNAscope technology **(Fig. 5a, 5b and Extended Data Fig. 6**). Similarly, *Pnoc* and *Grp* co-expression delineated the cl3 cluster from the *Pnoc*-only cl4 cluster (**Fig. 3i, j)**. Different combinations of neuropeptide expression were associated with known motor pool regulators.

**Fig. 6:**
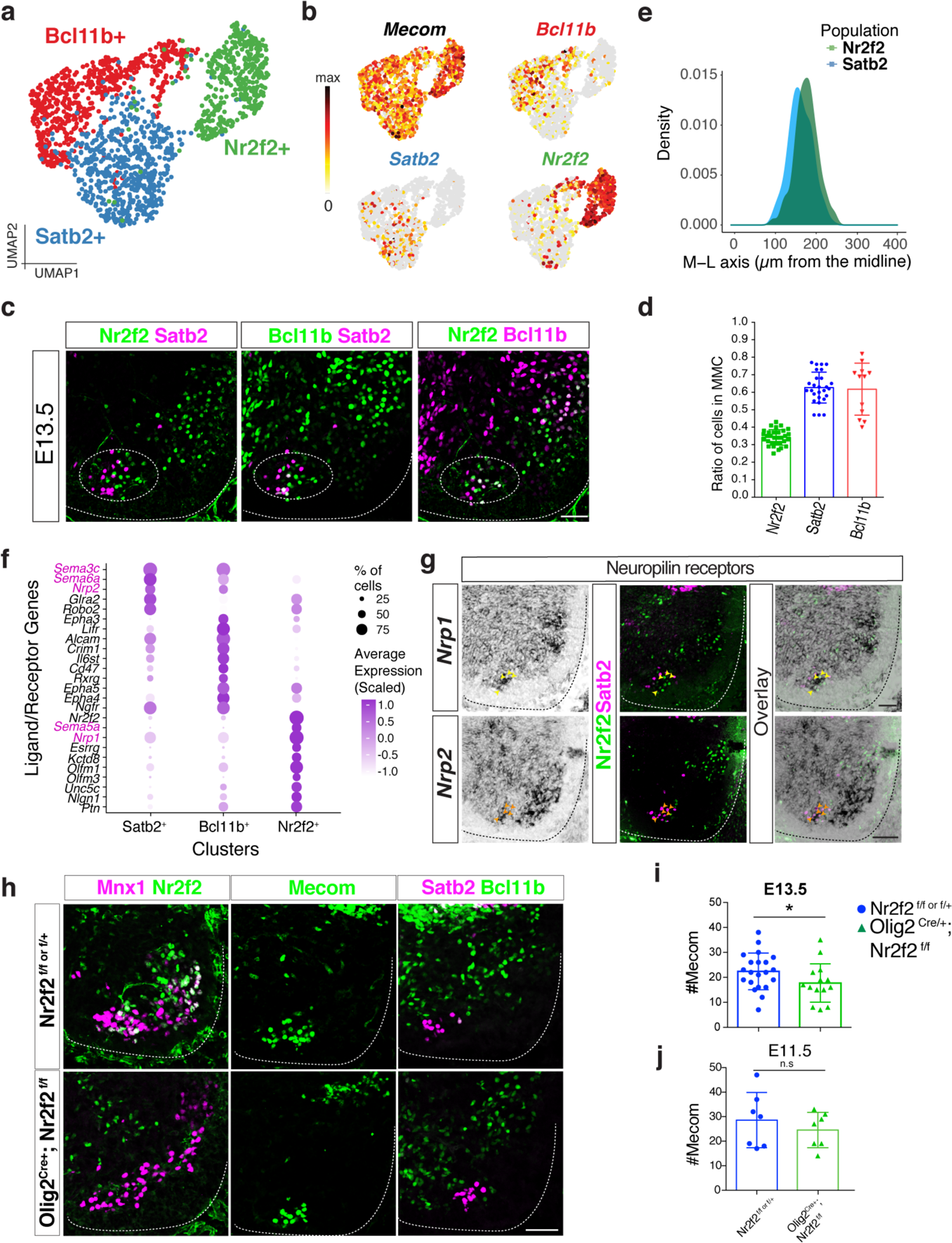
Molecular and functional heterogeneity in MMC neurons (a) UMAP visualization of brachial MMC cells with three molecularly distinct subpopulations. (b) Normalized expression of marker genes on UMAP space. *Mecom* is a known marker for MMC MNs and is highly expressed in all MMC neurons, but *Bcl11b*, *Satb2* and *Nr2f2* are enriched in different subpopulations. (c) Immunostaining for the MMC subpopulation markers Satb2, Nr2f2 and Bcl11b in brachial segments of E13.5 spinal cord. Nr2f2 and Satb2 are expressed in a mutually exclusive manner, whereas a fraction of expression of Bcl11b overlaps with both these latter. (d) Quantification of cell numbers in different subpopulations of Mecom*^+^* MMC neurons in brachial E13.5 spinal cord. Results are shown as mean ± SD, representing average counts from ≥3 sections of brachial spinal cord per embryos. For Nr2f2^+^, Satb2^+^ and Bcl11b^+^ population, n = 31, 25 and 12, respectively. (e) Mediolateral (M-L) density plot of the Satb2^+^ (blue) and Nr2f2^+^ (green) subpopulations in brachial E13.5 spinal cord. Data from 12 embryos. (f and g) Dot-plot showing selective expression of ligand/receptor genes in MMC subpopulations, revealing that *Nrp1* and *Nrp2* are differentially expressed in the *Nr2f2^+^* and *Satb2^+^* subpopulations, respectively (f). This result was validated *in vivo* by means of *in situ* hybridization (ISH) of *Nrp* genes and post-ISH immunostaining for Nr2f2 and Satb2 (g). Yellow and orange arrowheads indicate *Nrp1*:*Nr2f2* and *Nrp2*:*Satb2* co-expressing cells, respectively. (h and i) Genetic manipulation of Nr2f2 conditional knockout (cKO) from MNs by crossing Nr2f2 floxed mice with Olig2^Cre/+^ mice. Immunostaining for Nr2f2 and Mnx1 validated complete Nr2f2 loss from the MNs. Reduced numbers of Mecom*^+^* MMC MNs were apparent in the Nr2f2 cKO line (Olig2^Cre/+^; Nr2f2^f/f^) relative to control (Nr2f2^f/f or f/+^), with quantification shown in (i). Immunostaining for Satb2 and Bcl11b (quantification in Extended Data Fig. 9e) revealed no significant change in the ratio of these subpopulations. Results represent mean ± SD, quantified from n=9 and n=6 embryos for control and mutant mice respectively, with ≥2 brachial segments from each embryo. Statistical significance is represented as *p < 0.05, based on two tailed t-test. (a) (J) Quantification of Mecom*^+^* MMC neurons at E11.5. Results represent mean ± SD, quantified from n=4 and n=3 for control and mutant embryos, respectively. n.s denotes no significant difference based on two tailed t-test. Scale bar for all images in this Figure represents 50 μm.

For instance, Etv4^+^ motor pools (cm1) expressed *Sst*, whereas Pou3f1^+^ motor pools (cm3) displayed *Penk* expression (**Fig. 5c, d)**. Given that Hox cofactors and LIM homeodomain (HD) combinatorial codes govern distinct subsets of LMC MNs projecting along motor nerves^10, 46^, we propose that the battery of effector genes we have identified could be regulated by TF codes, and that the combinatorial expressions of receptors and ligands, cell adhesion molecules, and neuropeptides ensure precise neuromuscular and sensorimotor connections.

### Convergent and divergent clusters between brachial and lumbar LMC MNs

We also investigated putative subcluster identities in lumbar LMC MNs. We allocated the rostrocaudal and mediolateral positional information of “spatial quadrants” **(Extended Data Fig. 7a**) and performed subclustering within each spatial quadrant, leading to the identification of 10 subclusters with distinct molecular expression patterns **(Extended Data Fig. 7b-e, Supplementary Table 9**). Next, we probed if there is a lumbar LMC cluster analogous to the brachial LMC cluster. To effectively compare LMC subclusters between brachial and lumbar segments, we performed integration analysis of LMC cells from brachial and lumbar and jointly projected these cells onto a shared UMAP space using Harmony algorithm^59^. By examining the degree of cell group mixing across segments and performing quantitative similarity analyses based on the mutual nearest neighbors of each cell in the UMAP space, we were able to assess the relatedness of LMC cells across brachial and lumbar (**Fig. 5e-h**). First, we observed that brachial medial clusters are more similar to lumbar medial clusters and *vice versa*, suggesting that the molecular basis of mediolateral identity is shared by both segments (**Fig. 5f-h** and Extended Data Fig. 7f). Second, some known subclusters, including *Etv4*^+^, are found in both the brachial and lumbar segments **(Fig. 5i, left panel and Extended Data Fig. 8c, d)**. Third, there are LMC subclusters which appear to be segment-specific. For example, *Grp*^+^ cl3 and rl5 subcluster are present in brachial segments, whereas *Nkx6.1*^+^ crl6 subcluster predominantly occur in lumbar segments (**Fig. 5i**, middle and right panel, **Extended Data Fig. 8c, d)**.

**Fig. 7:**
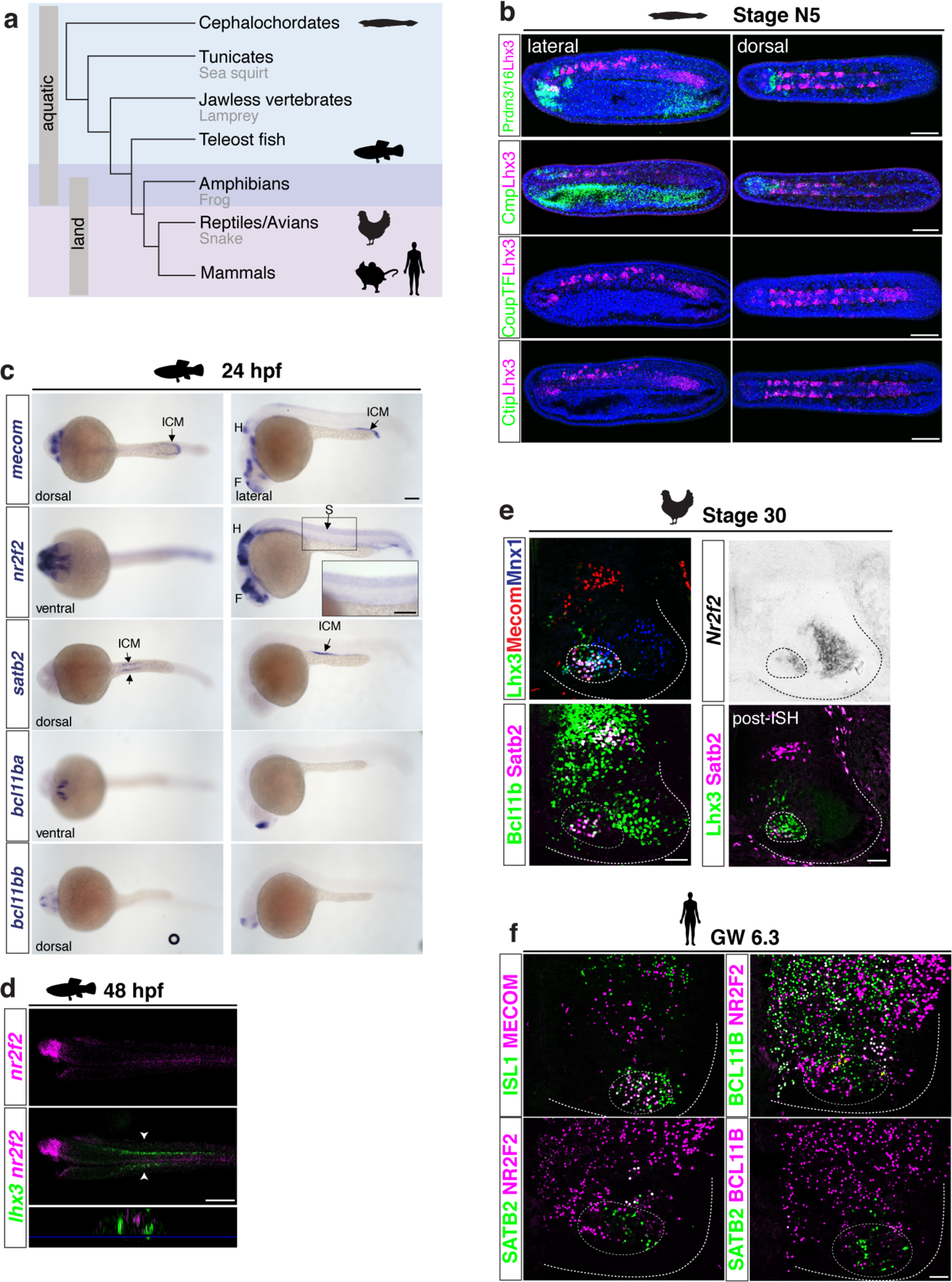
MMC diversification across chordate evolution. (a) Schematic illustration of chordate phylogeny, including marine invertebrate chordates (cephalochordate amphioxus and tunicates), aquatic vertebrates (jawless vertebrates and fishes), and land vertebrates (amphibians, birds and mammals). (b) Dual-FISH of amphioxus *Prdm3/16*, *Cmp*, *CoupTF*, and *Ctip* (homologs of rodent Mecom, Nr2f2 and Bcl11b, respectively) reveal that expression of none of these MMC subtype markers is detectable in the *Lhx3*^+^ cells of amphioxus nerve cord at stage N5. Left panels show lateral views of embryos, right panels show dorsal views (anterior to the left). Blue represents DAPI signal. Scale bar represents 50 μm. (c) In 24 h post-fertilization (hpf) zebrafish embryos, only *lhx3* and *nr2f2* were detected within the nerve cord, but not *mecom*, *satb2*, *bcl11ba* or *bcl11bb*. Zebrafish *bcl11ba* and *bcl11bb* are paralogs of the *Bcl11b* gene. A high-magnification image is presented of *nr2f2* expression within the somite region. All images on the left reflect dorsal views except for *nr2f2* and *bcl11ba* (ventral), and all panels on the right show lateral views. ICM: intermediate cell mass; S: spinal cord; H: hindbrain. Scale bar for all images represents 100 μm. (d) Dual-FISH of *nr2f2* and *lhx3* demonstrates that *nr2f2* expression is not localized in *lhx3^+^* MNs at 48 hpf. An orthogonal cross-section is shown in the bottom panel. Arrowheads indicate region where cross-section is performed. Scale bar represents 100 μm. (e) Immunostaining for Mecom, Lhx3 and Mnx1 (upper left) in Stage 30 chick spinal cord shows that all mature MMC MNs are Mecom*^+^*. Similar to mouse embryos (Fig. 6), ISH of *Nr2f2* and post-ISH immunostaining for Lhx3 and Satb2 (upper and bottom right) show mutual exclusive expression of *Satb2* and *Nr2f2* with preferential mediolateral distribution. Bottom left image shows that Bcl11b is co-expressed in some *Satb2^+^* MMC MNs. Dashed lines outline spinal cord boundaries and dashed circles demarcate MMC boundaries based on *Lhx3* expression. Scale bar represents 50 μm. (f) Immunostaining for ISL1 and MECOM verifies that all ventromedial MECOM^+^ cells in human gestation week (G.W.) 6.3 embryonic spinal cord are MNs. NR2F2, SATB2 and BCL11B are all expressed in human MMC neurons. Scale bar represents 50 μm.

Next, we explored which molecular signatures delineate the brachial and lumbar LMC MN subtypes. Rhythmic locomotor patterns in mammals rely on coordination of the neuronal network controlling the arms/forelimbs and legs/hindlimbs^60^. Although LMC MNs occur in both the forelimbs and hindlimbs, the convergent and divergent molecular identities of the motor pools within the fore/hindlimb LMC neurons remain enigmatic^3^. We compared gene expression in LMC MNs from both segments and uncovered 112 DEGs, with *Hox* genes predominating **(Extended Data Fig. 8a, Supplementary Table 10**). This outcome is consistent with previous studies showing that *Hox* genes are the crucial TFs defining segmental MN identities^61^. Additionally, gene ontology (GO) analysis demonstrated that other pathways—such as MN cell fate specification and development, insulin signaling pathway and translation regulation—were more prominent in lumbar relative to brachial LMC MNs **(Extended Data Fig. 8b**), corroborating the notion that the developmental process adopts a sequential rostral-to-caudal pattern^4, 62^. Notably, one neuropeptide-related GO term also emerged from the segmentally-enriched GO dataset **(Extended Data Fig. 8b**) We performed *in situ* hybridization and detected expression of the neuropeptides *Grp* and *Npy* in subsets of brachial LMC MNs, but less so in lumbar segments, validating our comparative analysis between brachial and lumbar LMC subclusters (**Fig. 5i, j)**. Conversely, *Igf1* was enriched in lumbar segments, and more specifically in *Hoxd10*^+^ medial LMC MNs. Thus, we have demonstrated that like TFs and cell adhesion molecules, neuropeptides not only exhibit strong heterogeneous expression among LMC subclusters but they also differ between the brachial and lumbar segments. Neuropeptides are widely known as neuromodulators affecting behavior and sensory processing in postnatal mice^63, 64^, and our data indicate they play unappreciated roles in embryonic development that warrant further investigation. We elaborate on the importance of differentially-expressed neuropeptides in developing limb MNs and their possible functional implications in our Discussion.

### Identification of MMC MN subtypes and molecular signatures responsible for subtype survival

Our identification of potentially novel motor pools and markers in LMC neurons further prompted us to assess molecular heterogeneity in axial mucle-innervating MMC MNs. MMC neurons that enable undulatory locomotion are regarded as the ancestral MNs, based on their occurrence in primitive aquatic vertebrates, such as lamprey^65^. Despite a lack of systematic characterization of axial motor pools, we hypothesized that, like LMC MNs, MMC neurons are also comprised of molecularly distinct subpopulations. Indeed, a recent study uncovered molecular diversity within MMC MNs, with ∼40% of MMC neurons expressing *Ebf2* and loss of this gene resulted in a reduction of differentiated axial MNs^66^. Here, we wanted to tackle three fundamental questions relating to MMC neurons. First, how many MMC subtypes are there in mice? Second, what is the mechanism underlying MMC heterogeneity? Third, and most pertinently, have MMC MN subtypes evolved to become more complex? To tackle these three questions, we subtracted MMC MNs from rostral samples and revealed three subpopulations among them, namely Nr2f2^+^ (Coup-TFII), Satb2^+^, and Bcl11b^+^ (Ctip2) (**Fig. 6a-c, quantification in Fig. 6d**). Notably, the Satb2^+^ subpopulation included an Ebf2^+^ subtype of MMC MNs reported previously^66^ **(Extended Data Fig. 9a, b)**. We observed that Satb2^+^ and Nr2f2^+^ MMC neurons were expressed at E13.5 in a mutually exclusive manner (**Fig. 6c and Extended Data Fig. 9c**). Moreover, some Bcl11b^+^ MMC MNs co-expressed Nr2f2 or Satb2 (∼15% each). Given that the LMC neurons in limb segments separate into mediolateral positions to innervate proximal and distal peripherals, we examined if the MMC subpopulations display spatial organization. A topographical analysis revealed a preferential medial localization for the Satb2^+^ subpopulation among brachial MMC neurons relative to the Nr2f2^+^ subpopulation (**Fig. 6e**). Subsequent analysis of DEGs among these three subpopulations revealed selective expression of receptor and semaphorin guidance molecules.

The Satb2^+^ and Bcl11b^+^ subpopulations expressed *Sema3c*, *Sema6a* and *Nrp2*, whereas the Nr2f2^+^ subpopulation was enriched for *Sema5a* and *Nrp1* expression (**Fig. 6f, g)**. This result indicates that, similar to LMC MNs, MMC neurons are also subdivided into medial and lateral positions, selectively expressing TFs and axon guidance molecules that might facilitate axon targeting fidelity, as demonstrated previously^67^. Thus, our data support the hypothesis that MMC MNs consist of three molecularly and spatially defined subpopulations marked by Satb2, Nr2f2 and/or Bcl11b expression, which might innervate different epaxial muscles^6^.

Given that Nr2f2, Satb2 and Bcl11b are known to facilitate brain region patterning^68, 69^, we tested if these TFs mediate MMC subtype diversification. As a proof of principle, we conditionally knocked out Nr2f2 from MN progenitors by using Olig2^Cre/+^, Nr2f2^flox/flox^ (cKO) and observed a ∼30% reduction in MMC MNs at E13.5 (**Fig. 6h, i)**, representing approximately the size of the *Nr2f2^+^* subpopulation in control mice (**Fig. 6d**). To examine the role of Nr2f2 in MMC neurons, we examined if this nuclear receptor acts in cell fate assignment, maintenance or survival. At E11.5, the MMC neuronal populations were comparable between control and cKO mice, indicating that specification and generation of the *Nr2f2^+^* subpopulation had not been affected by cKO (**Fig. 6j**). However, cleaved caspase-3 was more prominently expressed in the MMC neurons of E12.5 cKO embryos relative to control **(Extended Data Fig. 9d**). Notably, the ratio of *Satb2^+^* and *Bcl11b^+^* MMC MNs was not affected upon loss of Nr2f2 function, unlike the inhibitive interaction between Satb2 and Bcl11b in the brain cortex (**Fig. 6h, right and Extended Data Fig. 9e**)^69^. Taken together, these results indicate that Nr2f2 is indispensable for maintaining *Nr2f2^+^* MMC MN subpopulation.

### MMC subtypes during chordate evolution

Aquatic and terrestrial vertebrates display distinct rhythmic locomotion. Most fishes generate propulsive forces via rhythmic contraction of their axial muscles during swimming, whereas tetrapod employ their axial muscles for non-locomotory spinal alignment and breathing^6^ (**Fig. 7a**). To establish the mechanism underlying this distinct deployment of axial muscles necessitates cross-species comparison of axial neuronal subtypes and motor circuits. Given that it is still unknown when the molecular diversity of MMC MNs emerged during evolution, our discovery of novel MMC subpopulations provides a basis for establishing how MMC diversity could have evolved.

To examine if the axial MNs of early vertebrates display heterogeneity, first we investigated MN diversity in the cephalochordate amphioxus, an early-branching chordate that acts as a model for understanding the ancestral traits of vertebrates^70^. Previous studies have demonstrated several features of nervous system development shared by amphioxus and vertebrates. For instance, in both cases, the nervous system is patterned along the anteroposterior axis by *Hox* genes and along the dorsoventral axis by BMP/Wnt/Shh^71, 72^. Moreover, orthologs of vertebrate *Mnx1* and *Islet* genes are expressed by amphioxus MNs during development^73, 74^. We confirmed that *Lhx3* and *Mnxa/Mnxb* are expressed in amphioxus MNs as punctate tracts along the neural tube at stage N5 **(Extended Data Fig. 10a**)^75, 76^. However, we did not detect expression of *Prdm3/16*, a potential ortholog of *Mecom*, in amphioxus *Lhx3^+^* MNs (**Fig. 7b**). To evaluate molecular diversity of MMC MNs in amphioxus, we characterized the expression patterns of amphioxus orthologs of our newly-identified MMC subpopulation markers, *Satb2*, *Nr2f2* and *Bcl11b*. Phylogenetic analyses revealed that vertebrate *Satb2* is potentially derived from the *COMPASS (Cmp)* gene of the common ancestor of chordates^77^, while vertebrate *Nr2f2* and *Bcl11b* are homologs of *CoupTF* and *Ctip* in amphioxus. Dual fluorescence *in situ* hybridization (FISH) demonstrated that orthologs of these vertebrate MMC subpopulation markers are not expressed in *Lhx3^+^* MNs, though the telencephalon-like region of amphioxus displayed prominent *Cmp* expression. Therefore, amphioxus embryos do not appear to exhibit MMC subpopulation markers, implying that MMC MN diversity arose later during vertebrate evolution.

In zebrafish, a representative model of aquatic vertebrates, *lhx3* is expressed in all postmitotic MNs by 24 or 48 hours post-fertilization (hpf)^78^ **(Extended Data Fig. 10b**). From 24-72 hpf, we observed dynamic expression of *mecom, satb2* and the two paralogous genes of *Bcl11b*, i.e., *bcl11ba* and *bcl11bb*, in different regions of the CNS. However, except *nr2f2*, none of these genes was expressed in the spinal cord of zebrafish (**Fig. 7c, Extended Data Fig. 10b, c)**. Although *nr2f2* expression was detected in zebrafish spinal cord at 24 and 48 hpf, FISH indicated that *nr2f2* expression does not colocalize with *lhx3^+^* MNs (**Fig. 7d**).

Next, we examined expression of MMC marker genes in chicken spinal cord using immunostaining and ISH of chick spinal cord sections. We detected Mecom expression in a majority of Lhx3*^+^* MMC MNs. Moreover, chicken MMC MNs also expressed Nr2f2, Satb2 and Bcl11b, with Satb2 and Nr2f2 expression being mutually exclusive and displaying a preferential mediolateral distribution similar to the distribution of MMC subtypes observed for mouse embryos (**Fig. 7e**).

Finally, to determine if MMC MN diversity is conserved in human, we investigated the distributions of SATB2, NR2F2 and BCL11B in human embryo spinal cord sections at Carnegie stage 18, from Gestational Week (GW) 6.3 (equivalent to mouse E13.5 embryos when most MN columnar identities have been established)^79, 80^ (**Fig. 7f and Extended Data Fig. 10d**). We detected segregation of SATB2*^+^* and NR2F2*^+^* subpopulations in the MECOM^+^ MMC MNs, similar to the expression in chicken and mouse embryos. In contrast to what is observed in mouse embryos, only a few BCL11B*^+^* appeared to be co-expressed with NR2F2. At GW 10.3, SATB2 and NR2F2 expression remained segregated, indicating that the distinction between subpopulation persists in human embryo development (**Extended Data Fig. 10d**). Taken together, our analysis indicates that despite MMC MNs from cephalochordates to primates sharing expression of the same master regulator, Lhx3, MN diversification varies across species. We found that subpopulations of MMC MNs marked by expression of Satb2, Nr2f2 and Bcl11b are only present in terrestrial vertebrates and are absent from aquatic vertebrates. Accordingly, we speculate that the emergence of MMC MN diversity accompanied the vertebrate transition from aquatic to terrestrial lifestyles, allowing axial motor circuits to be rewired and enabling vertebrates to adapt to life out of water.

## Discussion

How many neurons are generated in the CNS? How is the diversity of neurons generated? Why are only particular types of neurons vulnerable to neurodegenerative disease? These are three of the most fundamental questions in neuroscience and they have been puzzling neuroscientists, developmental biologists, and clinicians for centuries. Traditionally, neuronal types are defined by their morphology, distribution, connectivity, behavior and electrophysiological properties^1^. Here, we addressed these challenges by detailed characterization of mouse spinal MNs through a massive parallel scRNA-seq analysis.

### Implications of scRNA transcriptomic analysis for spinal MNs

Although scRNA-seq promises comprehensive profiling of heterogeneous tissues, it often falls short in identifying rare cell types due to insufficient numbers of cell samples. As a result, enrichment steps before profiling are often required to better capture the diversity among a given cell type^18^. Our single-cell gene expression profiling enabled robust identification of MN subtypes. First, we utilized the Mnx1-GFP reporter and FACS to enrich for MN populations. To ensure that the stress of the dissociation process did not exclude vulnerable subpopulations from our analysis, we adjusted the protocol so that high-quality spinal MNs could be collected. This approach enabled finer resolution inspection of subtype diversity within each motor column and it potentially represents an optimal strategy for other tissues. Indeed, a previous study focused on collecting diverse cell types in the cervical and thoracic spinal cord at multiple time-points captured only ∼800 postmitotic MNs^2^. Low sample number limits both unsupervised discovery of cell types and estimates of gene expression. The refined enrichment protocol presented here allowed us to collect thousands of MNs so that MN columnar and motor pool heterogeneity during development could be investigated in detail.

### Exploring the diversity of spinal MNs

MNs innervating distinct muscles of the forelimb have been described, but the differences between these LMC MN populations remain largely unclear, apart from their genetic determinants. Our study complements and extends what is known about LMC MNs by describing combinations of transcription factors and neuropeptides that delineate their subpopulations, and we have validated these novel markers *in vivo*. In contrast, axial muscle-innervating motor pools are much fewer in number and thus are less accessible to manipulation. Accordingly, there is a paucity of information about the diversity within MMC MNs. Our single-cell analysis provided a novel way of addressing the underappreciated diversity of MMC MNs. Moreover, our systematic characterization of axial MNs in amphioxus, zebrafish, chicken, mouse and human embryos has provided insights into the evolutionary origin of the molecular diversity of spinal MNs. Therefore, here, we can present a detailed and diverse transcriptomic profile of spinal MNs, each represented by unique expression of a set of molecular codes corresponding to anatomically distributed subtypes. These molecular codes may serve not only as markers, but also provide insights into how precise axon targeting and neural circuit patterning is achieved. This principle seems to be applicable to other neuronal types^81^ and adult MNs^26^. Unexpectedly, either by means of trypsin- or papain-based disscociation approaches, we acquired equivalent numbers of MMC (2354 cells) and LMC (2248 cells) MNs, which is inconsistent with the population ratio of brachial LMC to MMC MNs (4:1) in E13.5 mouse embryos^82^. This scenario may reflect that LMC MNs are intrinstically vulnerable to shear stress, regardless of how mild the dissociation approach applied, and it is consistent with previous reports indicating that LMC MNs are more susceptible to degeneration in amyotrophic lateral sclerosis^83^. Additionally, we noticed that although the rostrocaudal identities of spinal cord cell types are determined and maintained by cross-regulation among Hox proteins^10^, our single-cell analysis revealed that Hox transcripts, such as *Hoxa5* and *Hoxc8*, can still be co-expressing within the same cell, despite at protein level, they are known to be segregated. This outcome is consistent with our previous studies showing that post-transcriptional mechanisms such as those exerted by microRNAs are indispensable for governing unambiguously spatiotemporal Hox protein expression and for defining rostrocaudal MN subtype identities in the spinal cord^84, 85^. Thus, although single-cell gene expression analysis facilitates identification of transcriptionally distinct known and novel cell types and states, integration of other modalities such as proteomics with our dataset would reveal more information about gene regulation during the cell type determination process.

From our scRNA-seq dataset, we discovered two previously uncharacterized columnar MNs, i.e., Calb1^+^ MNs in the brachial region and Nfib^+^Grm5^+^ MNs in the sacral segment. We have defined these two new types of neurons as MNs based on the facts that: (1) they co-express MN hallmark genes *Mnx1*, *Chat*, and *Slc18a3* (*VAChT*); and (2) both types extend their axons from the spinal cord to peripherals. Although Calb1 is a salient marker for V1 subtype Renshaw cells^86^, the presence of a small number of Calb1^+^ MNs among adult spinal MNs has been reported previously^87, 88^. In those studies, it was hypothesized that MNs are protected from glutamate-induced and calcium-mediated excitotoxicity via increased *Calb1* expression. Given that we observed persistent presence of Calb1^+^ MNs to the postnatal stage, it is tempting to examine if they are resistant to neurodegenerative disease. Moreover, neuronal expression of the metabotropic glutamate receptor Grm5 was shown previously to play pivotal roles in development of the mammalian CNS^89^. For example, *Grm5* expression prevented programmed cell death in *in vitro* cultures of rat cerebellar granule cells during the early maturation stage. Intriguingly, our novel sacral Nfib^+^Grm5^+^ MNs express Grm5, which is believed to exert neuroprotective effects on spinal MNs^90^. Given that sacral MNs are disease-resistant^91^, our newly identified Nfib^+^Grm5^+^ MNs may have implications for tackling MN disease. Thus, our identification of two previously uncharacterized Calb1^+^ and Nfib^+^Grm5^+^ neuronal types from our scRNA-seq dataset may shed light on “disease-resistant” MN subtypes. Future detailed functional characterizations of these two new columnar types and their peripheral innervating targets are clearly warranted.

### Diversity among LMC MNs

Despite ∼60 limb motor pools having been catalogued, fewer than 10 have been molecularly characterized^92, 93^. In this study, we describe the molecular codes for 26 LMC subclusters (16 and 10 for brachial and lumbar LMC MNs, respectively) that display unique combinatorial neuropeptide and TF expression profiles. Our findings support the terminal selector hypothesis that combinations of Hox/Lim homeodomain TFs might elicit a battery of subtype-specific genes, including ligands and receptors, cell adhesion molecules, and neurotransmitter-related genes, which would serve as regulators to robustly govern the fate and functions of subtypes^94–96^.

We have demonstrated herein that neuropeptides are expressed in an MN subtype-specific fashion. For example, *Penk* is specifically expressed in the Pou3f1^+^ motor pool that innervates the flexor carpi ulnaris muscle. Though the roles of neuropeptides in MN development remain unclear, their unexpected diversity in neurons could point to functions beyond acting as a general neurotransmitter in embryonic spinal MNs. Only a few neuropeptides in MNs have been assessed in previous studies, potentially serving as trophic factors to promote AchR accumulation and facilitating neuromuscular junction formation^97, 98^.

We hypothesize that these neuropeptides might: 1) modulate MN electrophysiology via autocrine signaling; 2) regulate motor circuit formation by guiding premotor interneuronal innervation to MNs; and/or 3) ensure the fidelity of finer innervation patterns to the peripherals. To better understand the functions of neuropeptides in neurons, it is tempting to investigate if motor pools display heterogeneous eletrophysiology patterns, if corresponding neuropeptide receptors are expressed on the innervated muscle subtypes, and if these trophic factors affect motor innervation patterns, such as singly- or multiply-innervated muscle fibers.

In relation to functional differences in terms of distinct wiring of motor circuits between rostrocaudal segments of the spinal cord, we anticipated that not all LMC subclusters would be conserved between brachial and lumbar segments. We identified some LMC subclusters presenting similar transcriptomic profiles (including of known markers such as *Etv4*) in both brachial and lumbar segments. However, some LMC subclusters sharing similar expression profiles expressed different known markers in different segments. For instance, *Pou3f1* is expressed only in brachial LMC MNs, but not in lumbar counterparts. Moreover, we also noted that some LMC clusters were only present in a specific segment. Thus, the differential neuropeptide enrichment among brachial and lumbar segments might provide insights into the potential differences in motor circuitry mediated by LMC motor pools.

While we made substantial progress in deciphering and characterizing LMC diversity compared to the observations in previous studies, our scRNA-seq approach did not characterize all ∼60 known LMC motor pools. There are two possible reasons for this outcome. First, the molecularly distinct identities of motor pools at the embryonic stage might not be reflected by the transcriptome alone. Increasing number of studies have demonstated that single-cell multi-omics analysis such as joint single-cell transcriptomic and epigenomic profiling likely increases the capability of dissecting cellular heterogeneity^99, 100^.

We envisage that multimodal analysis of spinal MNs would allow systematic positional allocations for all MNs, including our identified LMC subclusters and other vulnerable LMC neurons. Doing so may facilitate predictions of their innervating muscle targets^22^. Second, some neuronal identities may only be elicited after spinal cord wiring is complete at the postnatal stage. Accordingly, an examination of spinal MN identity at later time-points may prove enlightening^95^. Although we determined the number of subclusters using a robust ensemble approach based on spectral graph theory in mathematics, the inherent noisy and sparse nature of single-cell transcriptomics bring additional challenges to the estimation of optimal number of clusters in the data. Further computational subclustering analysis together with *in vivo* validation will likely increase our understanding of LMC diversity for more comprehensive identification of motor pools in LMC.

### MMC MN heterogeneity in chordates

Axial MNs that control core muscles represent an evolutionarily ancient neuronal type and they play pivotal functions across most vertebrate species. Although LMC neuronal diversity and respective limb motor circuits are relatively well characterized, diversity within the relatively small population of MMC neurons in the spinal cord is less appreciated^6^. In this study, we describe molecular heterogeneity within MMC MNs. In addition to characterizing the expression of distinct sets of molecular markers in MMC subpopulations, notably we observed mediolateral expression patterns for *Satb2* and *Nr2f2*. That finding prompts us to speculate that these MMC MNs might innervate different axial muscle groups, such as the longissimus, iliocostalis and the levator costae muscles^101^. Retrograde labeling and immunostaining of the MMC subpopulation markers identified herein will likely reveal the corresponding functional significance of MMC MN diversity.

Moreover, we have demonstrated that loss of the Nr2f2^+^ subpopulation led to a reduction in numbers of MMC MNs, but it did not affect the other two MMC subpopulations, i.e., Satb2^+^ and Bcl11b^+^. A previous study documented an evolutionarily conserved role for Nr2f2 in progenitor cells, acting as an important switching factor to regulate proper production of subtypes during the early-to-later born and neurogenic-to-gliogenic transitions^102^. Our study focused on postmitotic neurons and we report a potential role for Nr2f2 in maintaining a subset of MNs. In terms of the other two MMC subpopulation markers, *Satb2* and *Bcl11b*, they exhibit cross-repressive regulation in the vertebrate brain. Satb2 represses Bcl11b during specification of cortical L2-5^69^. Given that we also recently reported a miR-34/449/Satb2 regulatory axis in spinal interneurons^103^, we postulate that the microRNA-TF gene regulatory network could be involved in determining MMC subpopulation fate.

It remains largely unknown how axial-innervating motor circuits accommodated the change from undulatory locomotion to postural maintenance during the vertebrate transition from water to land^6, 104^. We hypothesized that rewiring of neural circuits accompanied gain or loss of neuron subtypes and so we investigated how MMC neuronal heterogeneity had evolved based on *Nr2f2/Satb2/Bcl11b* expression in amphioxus, zebrafish, chicken, mouse, and human embryos. We observed that Lhx3^+^ MMC neurons are present in all species examined, substantiating the notion that Lhx3^+^ MMC neurons represent the ancestral MNs^105^, which can be dated back to the common ancestor of all chordates. Although we did not detect *Satb2*, *Nr2f2* or *Bcl11b* expression in zebrafish MNs, different MN subtypes have been identified by previous studies^6^, such as the primary and secondary MNs born during two waves of neurogenesis that are distinguishable by cell body size and their fast and primarily slow muscle innervating types, respectively. We cannot completely rule out that other gene markers characterize some MMC subpopulations. For instance, generic MN genes, such as *mnx* paralogs, *islet1* and *islet2* are dynamically expressed in various combinations in different MN subtypes^106, 107^. Whether zebrafish embryos adopt newly evolved gene expression sets or simply utilize dynamically generic MN marker combinations to establish MMC subtypes remains to be established. A complete molecular profile of zebrafish MN subtypes at embryonic and adult stages would illuminate this topic more clearly.

Intriguingly, all three identified MMC subtypes are present in chicken, mouse and human embryos. Accordingly, we hypothesize that MMC diversification driven by Satb2/Nr2f2/Bcl11b might have allowed vertebrates to expand their MMC MN functions from locomotion to postural maintenance and spine alignment, facilitating more sophisticated limb muscle movement and thereby enabling the aquatic-to-terrestrial transition. Interestingly, a recent elegant study revealed that the neuronal subtype essential for walking originated in primitive jawed fish^108^, highlighting that the common ancestor of most vertebrates already possessed a sophisticated blueprint for walking. Here, we propose concordant neuronal rewiring during the aquatic-to-terrestrial transition. While the walking circuit posited from the tetrapod common ancestor existed ∼420 million years ago, the ancient undulatory MMC circuit diversified in parallel to accommodate an altered and more complex moving gait on land.

Overall, our study provides a unified classification system for deciphering the molecular repertoire of developing spinal MN and paves the way to establishing how axial and limb MN subtypes evolved. Our identification of combinatorial TF/neuropeptide markers for each MN subtype could also facilitate to identify differential susceptibility in disease models in search of disease-protective pathways that could be therapeutic targets.

## Extended Data Figures

**Extended Data Fig. 1:**
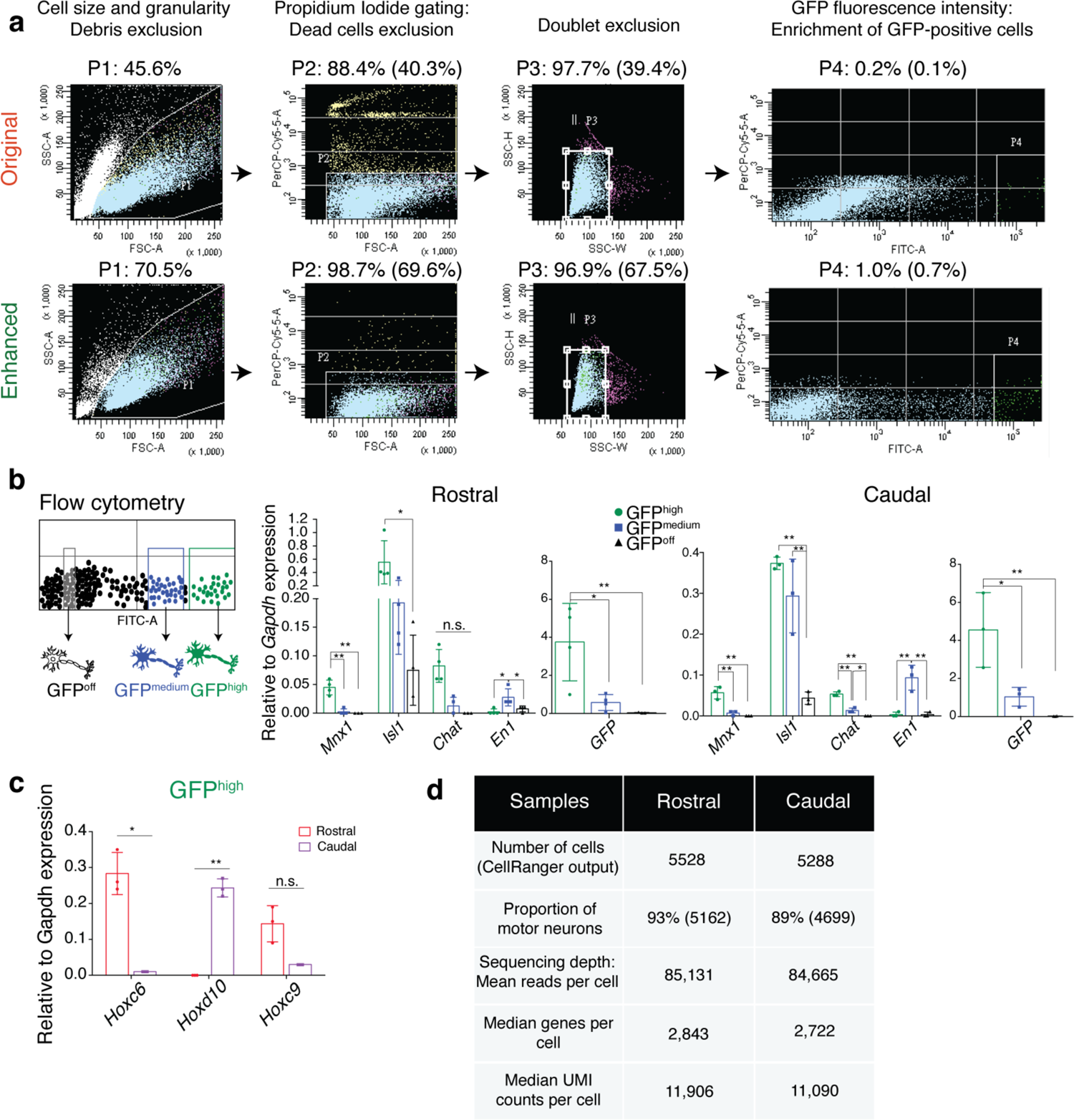
Optimization of single-cell dissociation of spinal motor neurons. (a) FACS plots comparing sample purity (P1), viability (P2), singlet or multiplets (P3), enrichment (P4) and yield before (upper) and after (bottom) optimization of MN dissociation protocols. The enhanced protocol was adapted for our single-cell RNA-sequencing experiment in this study. Percentages of desired populations are indicated above each plot, and total populations after subsequent gating are shown in parentheses. (b) Expression levels of MN markers and *GFP* genes relative to *Gapdh* from samples displaying differing GFP intensities (illustrated in the schematic at left), as measured by RT-qPCR (GFP^high^: green; GFP^medium^: blue; GFP^off^: black). Results are shown as mean ± SD from three independent experiments. (c) Expression levels of *Hox* genes to reflect segmental identities of the GFP^high^ sample. (d) Summary of median gene numbers, UMI counts, and cell numbers collected for each sample using our enhanced protocol. For B and C, statistical significance is represented as *p < 0.05, ** p < 0.01, n.s. = non-significant for one-way ANOVA with Tukey’s multiple comparison and paired t-test, respectively.

**Extended Data Fig. 2:**
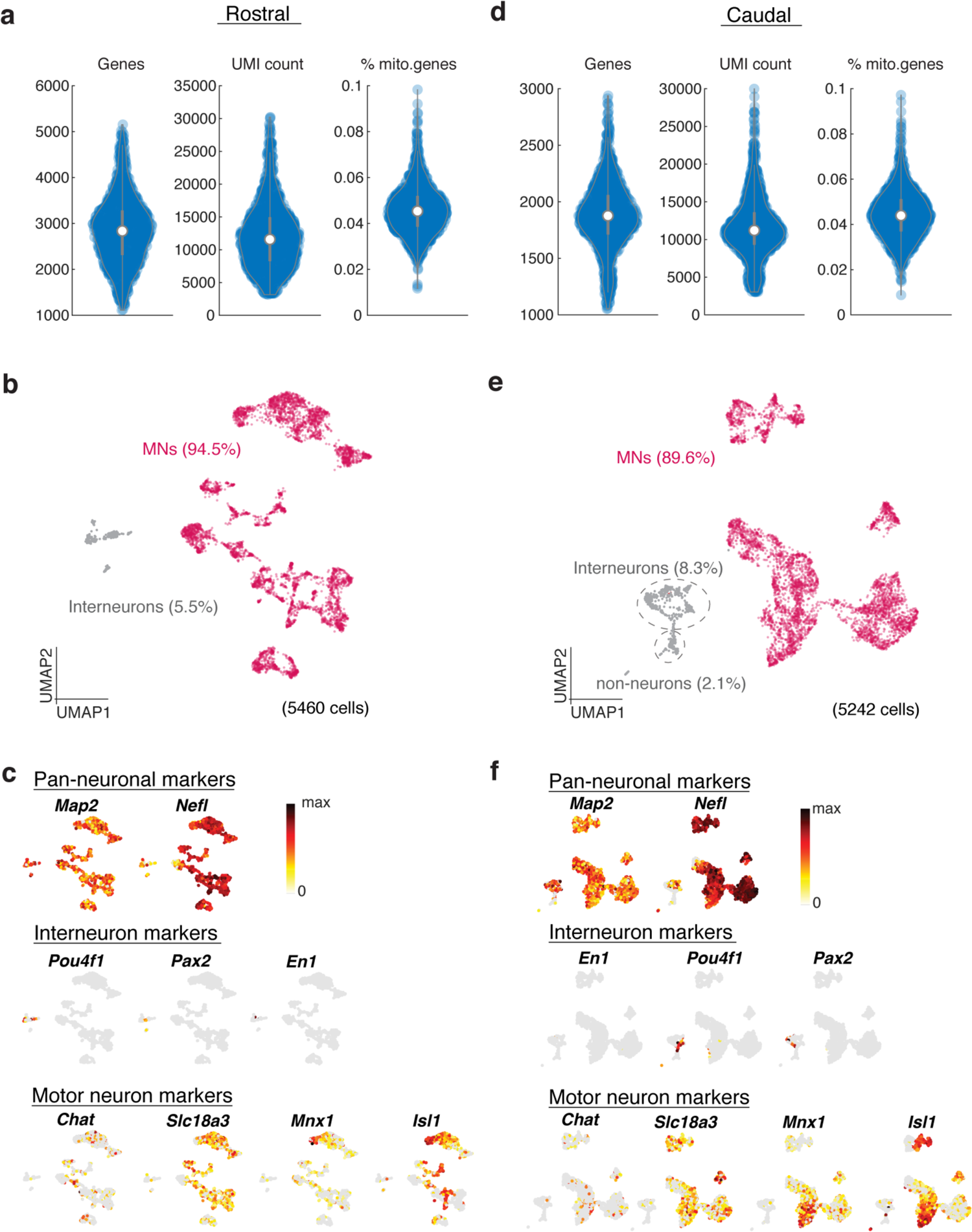
Quality control of collected samples. (a) Distribution of gene numbers and UMI counts, as well as the percentages of mitochondrial genes in the sequenced cells collected from rostral segments after removing low-quality cells. (b) UMAP visualization of all sequenced cells from a rostral sample. Cells are grouped into MNs (red) or interneurons and non-neurons (grey) based on expression patterns of known markers shown in (c). Percentages of each cell population in each sample are shown in parentheses. (b) UMAP distributions for expression in rostral samples of pan-neuronal, representative interneurons and MN markers. (d-f) As in (a-c), except caudal samples were analyzed.

**Extended Data Fig. 3:**
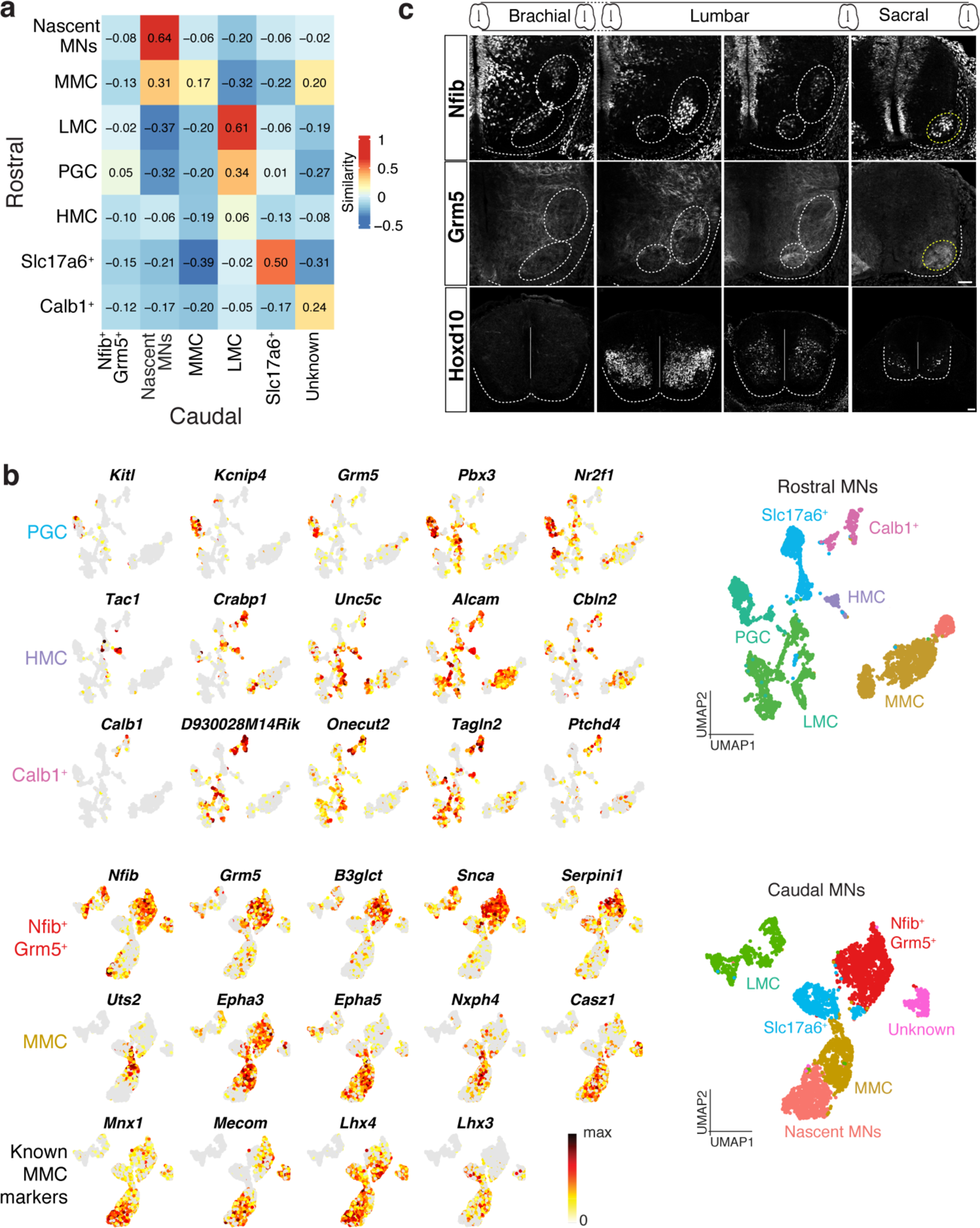
Identification of novel MN subtypes and marker genes for known and uncharacterized MN subtypes. (a) Similarity analysis between clusters from rostral and caudal segments based on the differentially expressed genes (DEGs). (b) UMAP visualization of the expression patterns of top marker genes for each cluster according to cluster identities shown in Fig. 1E and F (reproduced at right). MMC markers have been plotted to differentiate MMC MNs and *Nfib^+^* cells. (c) Immunostainings for Grm5 and Nfib across the rostrocaudal axis of E13.5 mouse spinal cord shows that double-positive *Nfib^+^Grm5^+^* cells are restricted solely to sacral segments. Hoxd10 immunostaining showed that its expression is only detectable in lumbar regions and diminished towards the posterior end of the spinal cord. Adjacent slides were used for Nfib and Grm5 immunostaining. Scale bar represents 50 μm.

**Extended Data Fig. 4:**
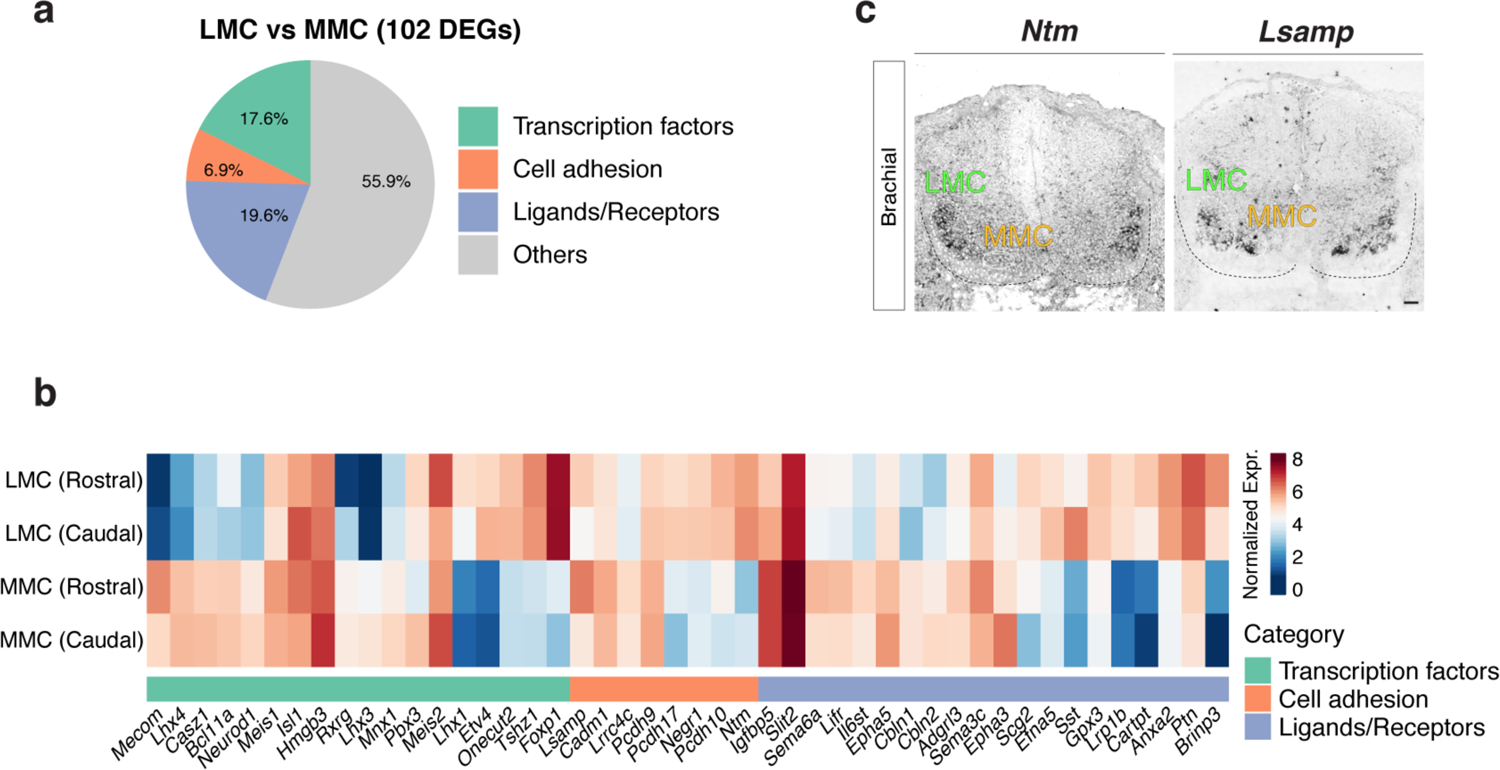
Differential gene expression between MMC and LMC MNs. (a) Percentages of TFs, cell adhesion molecules, ligand/receptors and others in the DEGs identified from MMC and LMC MNs. (b) Heatmap reflecting normalized expression of the top-ranked DEGs between LMC and MMC neurons in different categories. (c) ISH of the novel cell adhesion markers identified for LMC (*Ntm*) and MMC (*Lsamp*) MNs. Scale bar represents 50 μm.

**Extended Data Fig. 5:**
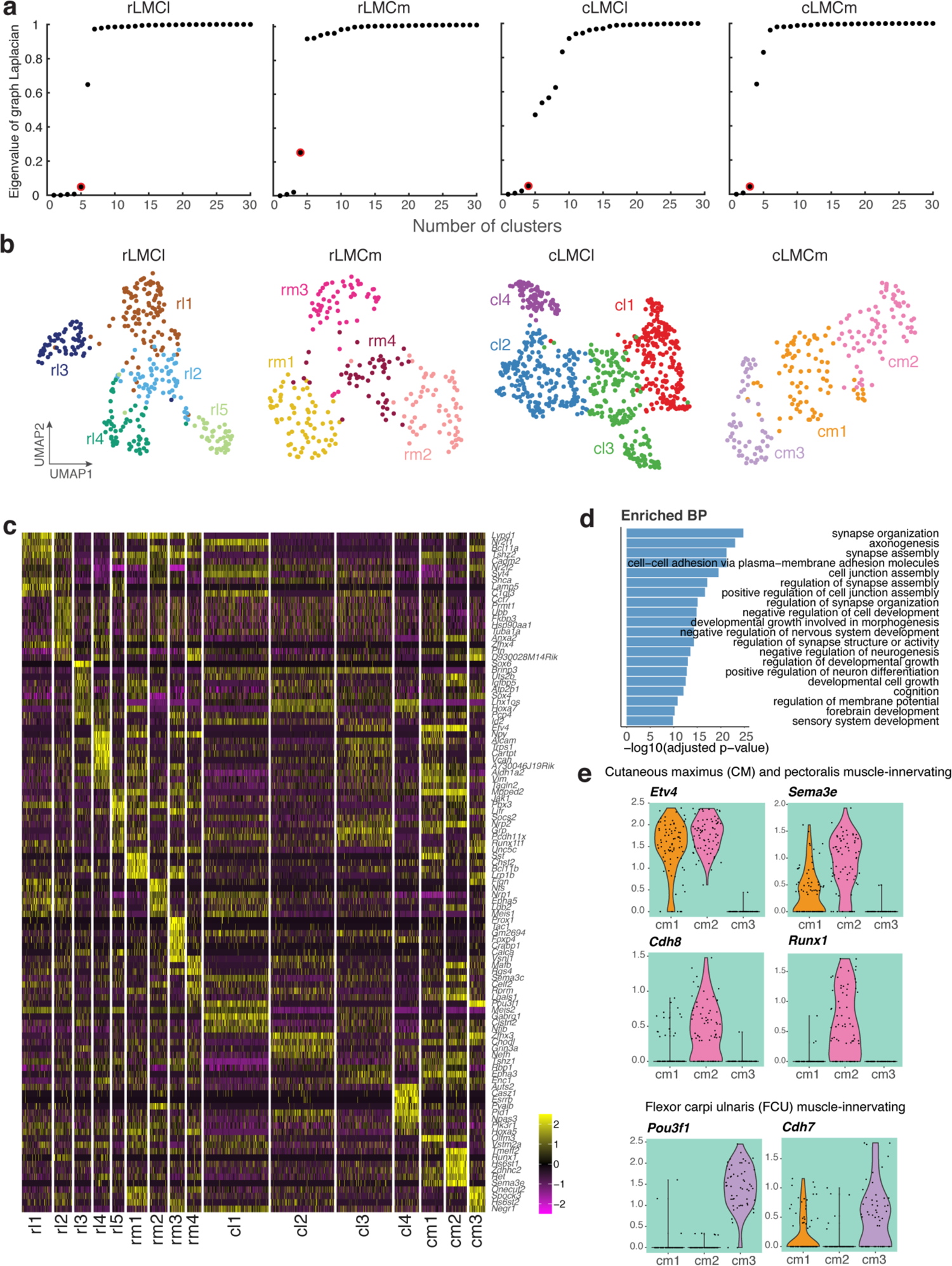
Sub-clustering analysis reveals heterogeneity among brachial LMC MNs. (a) Eigenvalue spectra reveal the number of subclusters in each “spatial quadrant”. The inferred number of clusters is marked in red, which is represented by the largest number before the largest gap of eigenvalue. r: rostral; c: caudal; m: medial; l: lateral. (b) UMAP visualization of cellular heterogeneity within each ‘spatial quadrant’ from brachial LMC MNs. Cells have been colored according to identified subclusters. (b) Heatmap showing scaled expression of the top 10 markers for each subcluster. (c) Gene ontology enrichment of biological processes of DEGs across brachial LMC neurons. Terms of interest in this study are highlighted in bold. (a) (e) Violin plots showing normalized expression of known markers. *Etv4*, *Sema3e*, *Cdh8* and *Runx1* are markers for motor pools innervating the cutaneous maximus and pectoralis muscles, whereas *Pou3f1* and *Cdh7* are markers for motor pools innervating the flexor carpi ulnaris muscle.

**Extended Data Fig. 6:**
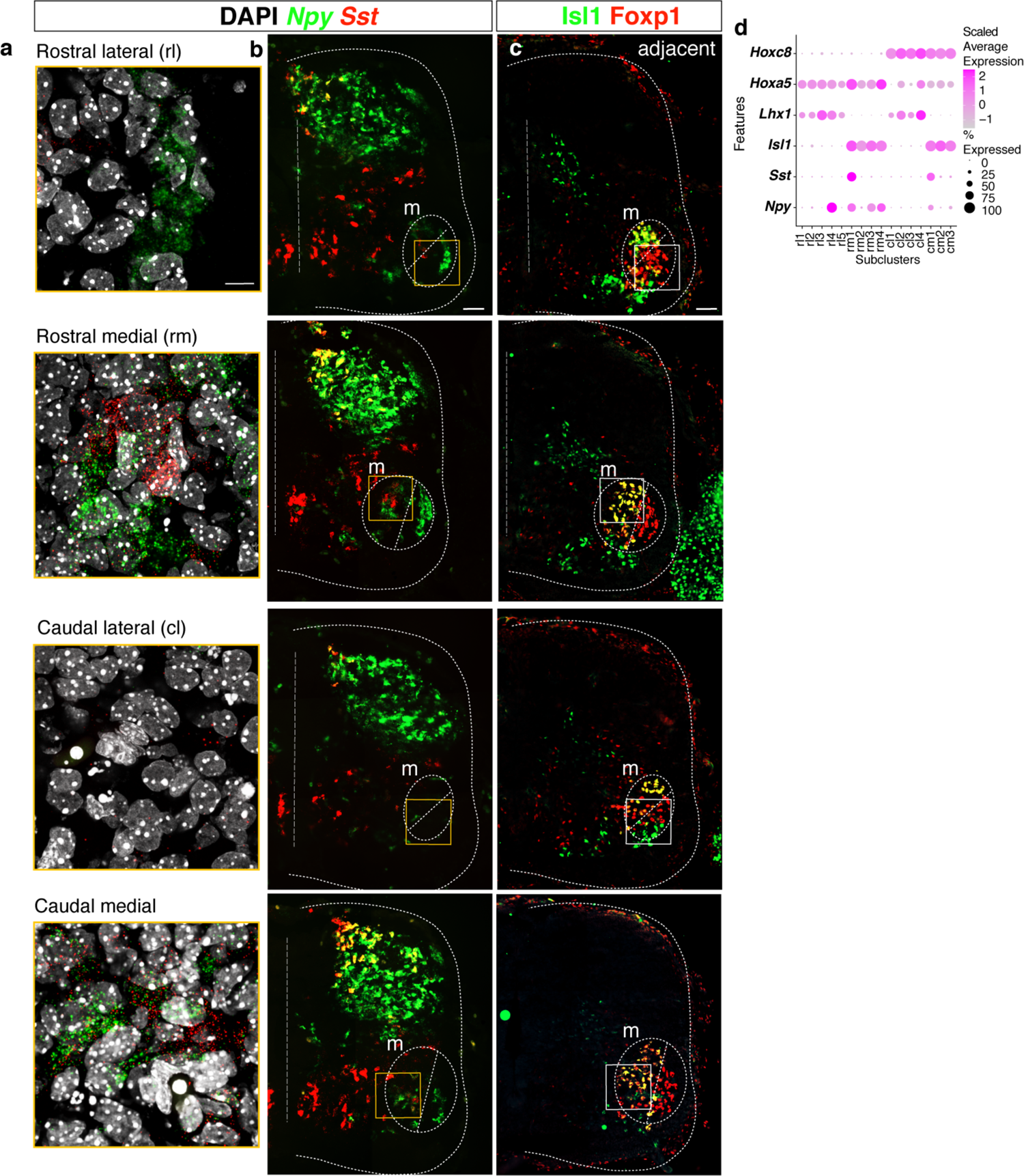
*Npy* and *Sst* display differential combinatorial expression in brachial LMC subclusters. (a) High-magnification images of *Npy* and *Sst* expression in LMC MNs (orange box in (b)), as detected using RNAscope technology. Representative images from each “spatial quadrant” are shown. Scale bar represents 10 μm. (b) Overview of the distribution of *Npy* and *Sst* marker expression in hemi-sections of the spinal cord. Scale bar represents 50 μm. (c) Staining for spatial markers on adjacent slides to (b) to delineate medial LMC (Foxp1^+^ Isl1^+^) and lateral LMC (Foxp1^+^ Isl1^-^) MNs. Scale bar represents 50 μm. (d) Dot-plot showing scaled average expression of *Npy*, *Sst* and spatial genes in each LMC subcluster. Circle size reflects the percentage of cells expressing the genes, and color intensity signifies the scaled average expression level of the genes. In b and c, white dashed contours outline the spinal cord boundary and spinal cord midline, dashed circles define LMC MN position, and the straight dashed white lines within the LMC demarcate the medial from lateral LMC MNs (m = medial). Orange boxes are shown as high-magnification images in (a), and white dashed boxes represent adjacent regions.

**Extended Data Fig. 7:**
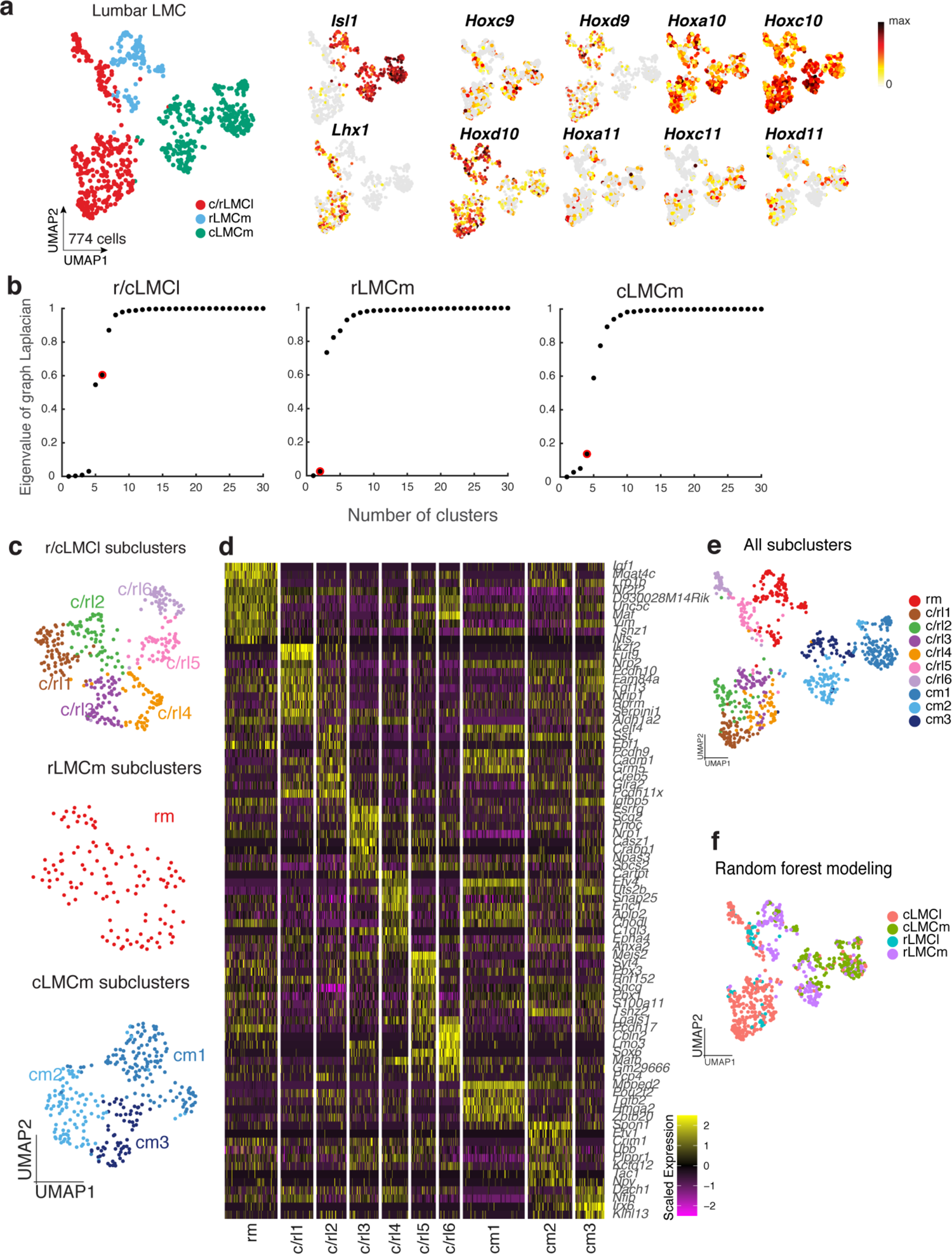
Sub-clustering analysis reveals heterogeneity among lumbar LMC MNs. (a) UMAP visualization of all lumbar LMC MNs. Cells have been grouped into “spatial quadrants” based on expression of *Hox* and LIM homeodomain genes. Note that among lateral LMC MNs, cells could not be separated into caudal and rostral lumbar segments. r: rostral; c: caudal; m: medial; l: lateral. (b) Eigenvalue spectra reveal the number of subclusters in each “spatial quadrant”. The inferred number of clusters is marked in red. (c) UMAP visualization of cellular heterogeneity within each ‘spatial quadrant’ of lumbar LMC MNs. Cells have been colored according to identified subclusters. (d) Heatmap demonstrating scaled expression of the top 10 markers for each subcluster. (e) UMAP distribution of all LMC subclusters. Cells have been color-coded according to the clustering results in (c). (f) Random forest modeling to predict cell identities of lumbar LMC MNs using the highly variable genes and cell identities from brachial LMC MNs. Similar to our unbiased clustering result shown in (a), this approach enables segregation between lateral and medial LMC MNs, but fails to distinguish rostral and caudal cell identities.

**Extended Data Fig. 8:**
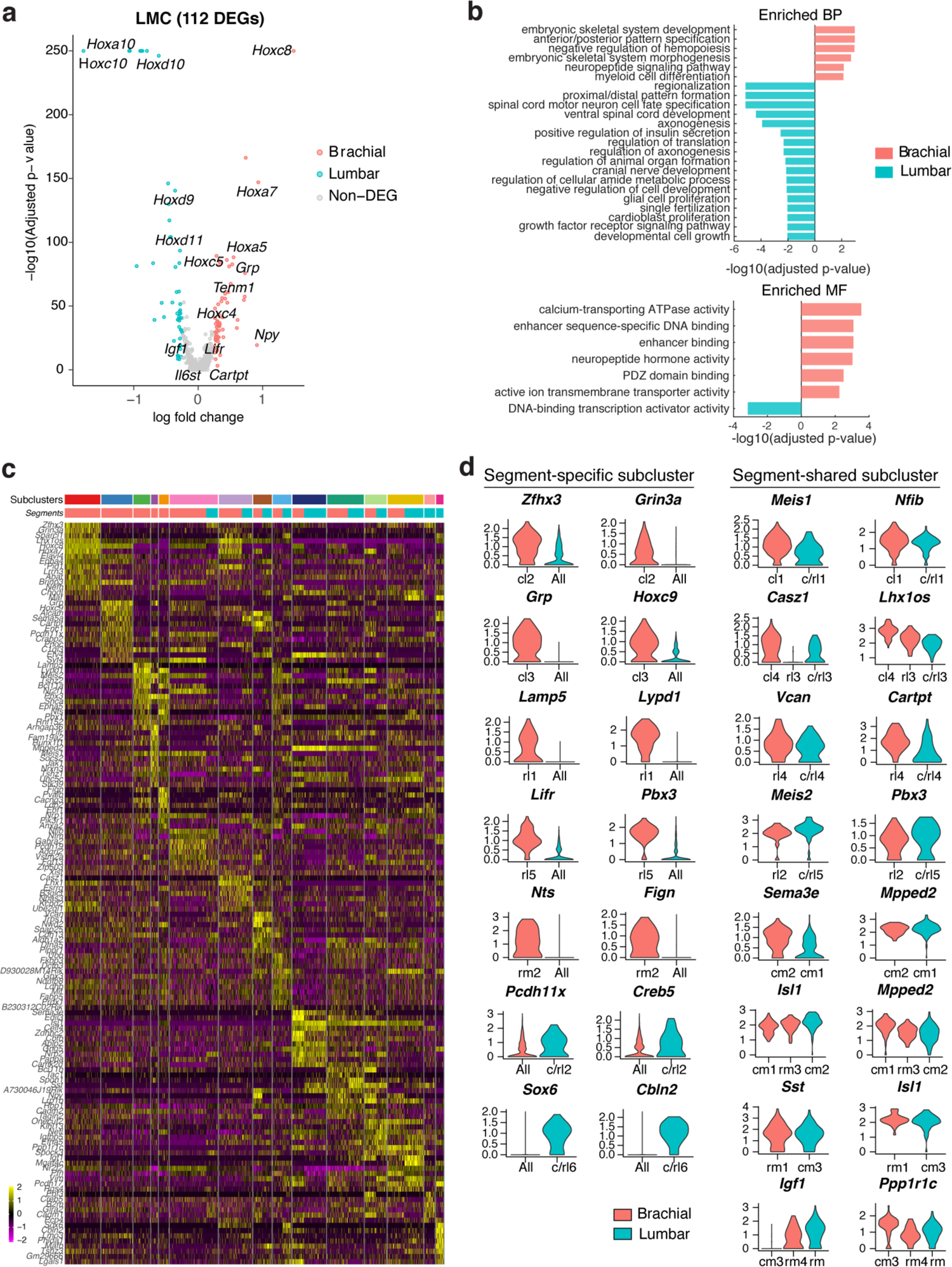
Comparison of LMC subclusters from brachial and lumbar segments reveals differentially expressed neuropeptides and *Hox* genes. (a) Volcano plot showing the DEGs between segments for LMC MNs. Colored dots reflect genes that exhibit significantly different expression. Genes with an adjusted p-value <0.05 and log fold-change >0.25 were deemed differentially expressed. The log-transformed fold-change is shown on the x-axis. *Hox* genes are the top-ranked markers differentially expressed between brachial (red) and lumbar (blue) segments. Note that a few neuropeptide genes (e.g., *Igf1*, *Npy*, *Grp*) were observed in the list of DEGs. (b) Enrichment analysis of DEGs between brachial and lumbar LMC neurons for biological process (BP, upper) and molecular functions (MF, bottom). Bold text highlights terms of interest in this study. Notably, the neuropeptide signaling pathway is shown as enriched in brachial segments. (c) Heatmap illustrating scaled expression of the top-ranked marker genes (rows) for cells in each group (columns) after merging brachial and lumbar LMC subclusters. Top (‘Subclusters’) bar labels subclusters from merged dataset. Second ‘Segments’ bar identifies cells from brachial (orange) and lumbar (turquoise) samples. (d) Violin plot shows expression of top two marker genes from each merged group (brachial, orange and lumbar, turquoise). Left panel displays gene expression in the segment-specific subcluster against all cells (All) from brachial or lumbar segments, while right panel shows gene expression from segment-shared subclusters.

**Extended Data Fig. 9:**
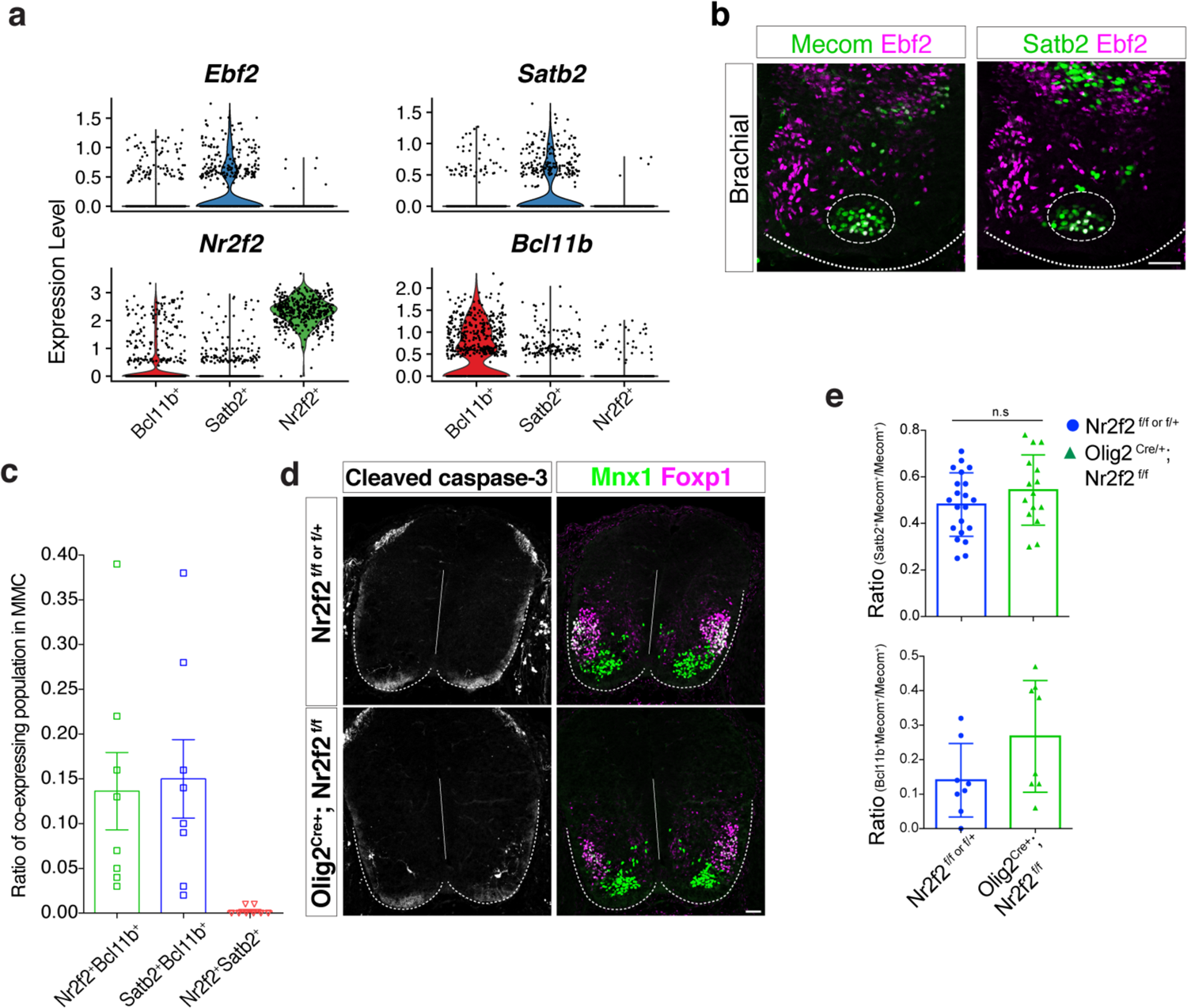
Molecular heterogeneity among MMC neurons. (a and b) scRNA gene expression (a) and immunostainings of Ebf2 and Satb2 (b) reveal co-expression of *Ebf2* in a fraction of the *Satb2^+^* population. Scale bar represents 50 μm. (a) (c) Quantification of the ratios of Bcl11b^+^:Satb2^+^, Bcl11b^+^:Nr2f2^+^, or Nr2f2^+^:Satb2^+^ co-expressing cells to total cells in MMC region of the spinal cord. Results are shown as mean ± SD, representing average counts from n=8 embryos with ≥2 sections of brachial spinal cord per embryo. (b) (d) Immunostaining for cleaved caspase-3 (an apoptotic marker), together with Mnx1 and Foxp1 to reflect motor column positions, in E12.5 control and Nr2f2 conditional knockout brachial spinal cords. Scale bar represents 50 μm. (c) (e) Quantifications of the Satb2^+^ and Bcl11b^+^ subpopulations upon Nr2f2 conditional knockout. No significant changes were observed. n.s denotes no significant difference based on two tailed t-test. Results represent mean ± SD. For Satb2^+^ ratio quantification, n=9 and n=6 embryos for control and mutant mice respectively, while n=4 each for both genotypes during Bcl11b^+^ ratio quantification. All embryos were investigated with ≥2 brachial segments.

**Extended Data Fig. 10:**
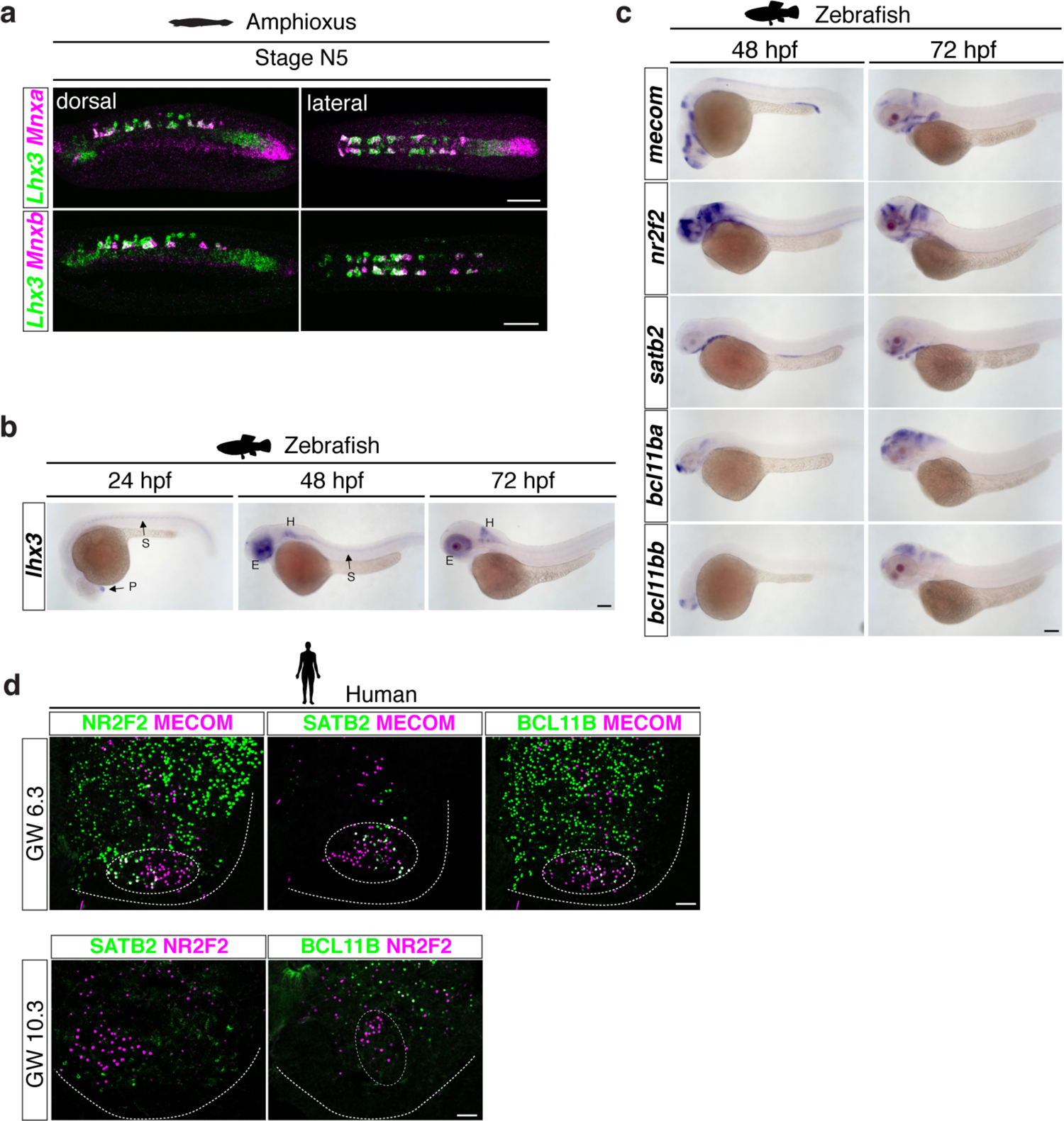
MMC diversity across chordate evolution. (a) Dual-FISH of *Lhx3* and *Mnxa*/*Mnxb* reveals punctate tracts in the nerve cord of amphioxus. Scale bar represents 50 μm. (b) ISH of *Lhx3* from 24 h post-fertilization (hpf) to 72 hpf reveal expression at both 24 and 48 hpf in the spinal cord of zebrafish. S: spinal cord; H: hindbrain; E: eye; P: pituitary gland. Lateral views of the embryos are presented (anterior to the left). Scale bar represents 100 μm. (c) ISH of MMC subpopulation markers in zebrafish embryos at 48 hpf and 72 hpf. Lateral views of the embryos are presented (anterior to the left). Scale bar represents 100 μm. (d) Immunostaining for the MMC subpopulation markers SATB2, BCL11B, NR2F2 and MECOM reveal they are expressed in subsets of the MMC MNs of human embryos at gestation week (G.W.) 6.3 and 10.3. Scale bar represents 50 μm.

## Methods

### Mice

Wildtype C57BL/6J, Mnx1-GFP^109^, Mnx1-RFP^83^; Calb1^Cre/Cre^ (JAX, 028532), Olig2^Cre/+^^110^, Tau-mGFP^111^ (JAX, 021162), and Nr2f2^f/f^^112^ mice were used in the study. During experiments, mice were crossed to obtain embryos of desired genotypes. When a copulation plug was observed, the embryo stage was estimated as E0.5. All of the live animals were maintained in a C57BL/6J background and housed in a specific-pathogen free (SPF) animal facility, approved and overseen by IACUC Academia Sinica guidelines.

### Sample collection for scRNA-seq

For single-cell spinal motor neuron collection, Mnx1-GFP mice were crossed to C57BL/6J mice. E13.5 embryos of pregnant mice were decapitated and dissected to isolate spinal cords from rostral (C4-T3) and caudal (L1-S5) segments in Leibovitz’s (L-15) medium. Tissue dissociation was performed using Neural Tissue Dissociation Kit (P) (Miltenyi Biotec, 130-092-628) on a gentleMACS dissociator (Miltenyi Biotec, 130-093-235) according to the manufacturer’s instructions. Dissociated cells were resuspended in N2B27/DMEM-F12 and neurobasal medium containing N2 (Life Technologies, 17502048) and B27 (Life Technologies, 17504044), 1% penicillin-streptomycin, 2 mM L-glutamine, 0.2 M ß-mercaptoethanol and 0.5 μM ascorbic acid, supplemented with 1% inactivated fetal bovine serum (FBS), 1:10000 DNase-I (Worthington Biochemical, LS006342), before filtering through a 70-μm strainer (Falcon, 352350). Sorting was carried out using a BDFACSAria III cell sorter (BD BioSciences, USA), with 85 μm nozzle diameter and 45 sheath pressure, to collect GFP^+^ cells at 4 °C into DMEM medium with 1% FBS. Collected cells were counted and adjusted to a final concentration of 700-1200 cells/μl.

### Single-cell library generation

ScRNA libraries were generated using a 10x Genomics Chromium Controller Instrument (10x Genomics, Pleasanton, CA) and Chromium Single Cell 3’ Reagent Kit v2 according to the manufacturers’ instructions. In brief, single-cell suspensions were loaded on the Chromium Controller Instrument to generate single-cell Gel Bead-In-Emulsions (GEMs). Upon breaking up of GEMs, the barcoded cDNA was purified and amplified. During library construction, the amplified barcoded cDNA was fragmented, A-tailed, and then adaptor-ligated. A sample index was added. Pooled libraries were sequenced using Illumina NextSeq 500 and NovaSeq S4 systems, and then paired-ended (read 1: 28 base pairs, read 2: 91 bp) to an average depth of 85k mean reads per cell.

### Preprocessing and quality control of scRNA-seq data

The FASTQ files were processed using the standard Cell Ranger pipeline (version 2.1.1, 10x Genomics) for demultiplexing, mapping to the mm10 reference, filtering, barcoding and to count unique molecular identifiers (UMI). For both rostral and caudal samples, cells were retained for subsequent analysis if they displayed a number of genes between 1000 and 5300, UMI counts <30500, as well as <10% mitochondrial counts.

### Clustering analysis of scRNA-seq data

We performed normalization, dimensionality reduction and cell clustering using Seurat package version 2.3.4^30^. The digital data matrices were normalized by a global method, whereby the expression value of each gene was divided by the total expression in each cell and multiplied by a scale factor (10,000 by default). These values were then log-transformed with a pseudocount of 1. Highly variable genes across cells were selected using the FindVariableGenes function (parameters: x.low.cutoff = 0.0125, x.high.cutoff = 3, y.cutoff = 0.5). To identify cell clusters, principal component (PC) analysis was first performed using the top 34 PCs for the rostral sample and the top 31 PCs for the caudal sample, where the number of significant PCs was determined based on an elbow plot and jackstraw test. Cells were then clustered using the Louvain community dection method in the FindClusters function (resolution = 0.05) and visualized using Uniform Manifold Approximation and Projection (UMAP). Motor neurons (MNs) and non-motor neurons were distinguished based on expression of generic MN markers such as *Mnx1* and the cholinergic genes *Chat*, *Slc18a3* and *Slc5a7*. Only clusters expressing MN markers were considered and further analyzed using the same pre-processing procedures as described above, but with different numbers of PCs and cluster resolutions.

To cluster MNs, cell clusters were obtained using the top 31 PCs with a resoultion of 0.08 for the rostral sample and the top 24 PCs with a resolution of 0.3 for the caudal sample. Cell identities were assigned using known markers established in previous studies, as well as the novel markers identified in our study. Marker genes of each cell cluster were identified using the FindAllMarkers function. Genes were considered marker genes if: (i) the adjusted p-values from the likelihood-ratio test was <0.05; (ii) the log fold-change was >0.25; and (iii) the percentage of cells expressing a given gene in the cluster was >25%.

To subcluster LMC neurons, cells were first grouped into “spatial quadrants” based on their expression pattern of *Hox* genes and LIM homeodomain factors. Cells in each “spatial quadrant” were then further clustered. In brief, for brachial LMC MNs, “spatial quadrants” were obtained by performing hierarchical clustering using *Hox* genes (*Hoxa5* and *Hoxc8*) and LIM homeodomain factors (*Lhx1* and *Isl1*). The robustness of assigned identities was confirmed by observing that the cells of a cluster within each “spatial quadrant” segregated from cells of a different cluster in UMAP space, which was established from PC analysis using highly variable genes as input. For the lumbar LMC MNs, “spatial quadrant” clusters were obtained by using the FindClusters function with a resolution of 0.3. The robustness of the assigned identities was confirmed by predicting cell identities among lumbar LMC MNs from the assigned identities of brachial LMC MNs using a random forest model. This was achieved by training a random forest classifier based on the “spatial quadrant” identities of brachial LMC MNs using the ClassifyCells function in Seurat, which was subsequently applied to the lumbar LMC MNs.

To further subcluster LMC MNs in each “spatial quadrant” cluster, we adopted a multi-resolution ensemble method to determine the number of subclusters based on spectral graph theory^113^. Specifically, we first performed subclustering using the FindClusters function in Seurat with multiple resolutions ranging from 0.1 to 3 and an increment of 0.05. Next, we constructed a consensus matrix based on a set of clustering results from these multiple resolutions. Consensus matrix *A* was obtained by averaging the entries of a connectivity matrix *B* across all runs, where the entry *B_ij_* of the connectivity matrix *B* equals 1 if cells *i* and *j* are assigned to the same cluster for each run. Thus, each entry *A_ij_* for *A* varies from 0 to 1 and represents the probability of cells *i* and *j* being in the same cluster across multiple resolutions. The consensus matrix was further pruned by setting the elements to zero if they were <0.3 to ensure better robustness against noise. Finally, we estimated the number of clusters by computing the eigenvalues of the associated Laplacian matrix of the constructed consensus matrix. Based on perturbation theory and spectral graph theory, it has been theoretically proven that the number of clusters *N* equals the multiplicity of the eigenvalue 0 of the Laplacian matrix ^113^. Therefore, in the ideal case of *N* completely disconnected clusters, the eigenvalue 0 has multiplicity *N*. More generally, the number of clusters *N* is usually given by the value of *N* that maximizes the eigenvalue gap (difference between consecutive eigenvalues), i.e., select the number *N* such that all eigenvalues are very small, but is relatively large. Once the number of clusters was determined, we ran FindClusters again to obtain the final clusters. In addition to this *in silico* inference, we also examined the heatmap of the top 10 marker genes of the identified clusters and ensured that the marker genes were biologically meaningful and that different clusters exhibited distinct expression patterns of those markers.

MMC MN subclusters were obtained by using the top nine PCs with a resoultion of 0.2.

### Similarity analysis between two groups of cells

Similarity between two groups of cells representing rostral or caudal MNs was determined using the calculateDistMat function (method = “trend”) in the CIDER package^114^. In brief, differentially expressed signatures (DES) were first identified using limma-trend data for each group of cells against all other cells within the dataset, whereby DES were computed by fitting a linear regression model. The similarity was then measured according to the Pearson’s correlation coefficient of DES between two groups.

To visually examine relateness among LMC subclusters of brachial and lumbar segments, we merged all LMC neurons from the brachial and lumbar segments and then performed joint dimensional reduction via UMAP by using the corrected embedding learned from the Harmony integration method^59^. To further quantify similarity among brachial and lumbar LMC subclusters, we assessed the degree of subcluster mixing across segments, which was quantified based on the overlap between one subcluster from brachial segments and another subcluster from lumbar segments in the joint UMAP space. Specifically, we first identified mutual nearest neighbors (MNNs) of each cell based on Euclidean distances in the UMAP space. Second, for each cell in one sample (e.g., brachial), we counted how many of its MNNs are cells in a certain subcluster from another sample (e.g., lumbar). Third, for each subcluster in the brachial sample, we computed its overlap ratio with another subcluster in the lumbar sample by calculating its average MNNs for cells from the lumbar sample, representing the probability of cells from two subclusters being MNNs across brachial and lumbar segments. Finally, to increase the robustness of the estimated similarity, we computed an average overlap ratio by using different numbers of neighbors (*k* = 10, 15, 20) to identify MNNs. The similarity matrix was visualized using the ComplexHeatmap R package^115^. To intuitively visualize the relationships of LMC subclusters between brachial and lumbar segments, we created a alluvial diagram using the ggalluvial R package (https://www.rdocumentation.org/packages/ggalluvial/versions/0.12.3). Lines of low similarity values (i.e., <0.09) were omitted to accentuate relatively strong similarity and enhance readability.

### Differential expression analysis and gene set enrichment analysis

Differentially expressed genes (DEGs) between segments or subclusters were identified using the FindMarkers function by performing likelihood-ratio tests. Genes with an adjusted p-value <0.05 and log fold-change >0.25 were considered differentially expressed. Gene Ontology (GO) enrichment analysis of DEGs was performed using the clusterProfiler R package (v3.14.3)^116^, focusing on Biological Processes (BP) and Molecular Functions (MF).

Redundant GO terms were removed using the *simplify* function with default parameters.

### Quantitative real-time PCR (qPCR)

FACS samples were lysed and extracted for RNA with a Quick-RNA MicroPrep Kit (Zymo Research). Total RNA (50-200 ng) from each sample was reverse-transcribed using Superscript III (Invitrogen) and one-tenth of the reaction product was used for qPCR. qPCR analysis was performed with technical duplicates using SYBR Green PCR mix (Roche) for genes of interest and *Gapdh* (a normalization control for amplification efficiency) on a LightCycler 480 Real-Time PCR machine (Roche). The primer sequence used for qPCR analysis is listed in Supplementary Table 12. Three independent experimental samples were analyzed.

### Primary MN culture, collection and immunostaining

Dissociated MNs from spinal cord were cultured on 0.01% Poly-L-Ornithine (Sigma-Aldrich/Merck, A-004-C) and 5 μg/ml laminin-coated (Thermo Fisher Scientific, 23017015) four-well plates with coverslips. Primary MNs were grown in medium containing Advanced DMEM/F12 (Invitrogen, 12634-010), 1% penicillin-streptomycin, 2 mM L-glutamine, B27, N2, and 0.1 ng/ml each of GDNF and BDNF for 48 h before harvesting and immunostaining.

To collect samples for immunostaining, similar procedures were conducted as described below for mouse embryo immunostaining, except that cells were fixed with 4% paraformaldehyde (PFA) for 15 min at room temperature. Samples were mounted with Aqua-Poly/Mount (Polysciences Inc.) and subjected to image acquistion using an ImageXpress® Micro XLS High-Content Imaging System (Molecular Devices). Cells stained with GFP and MN markers were quantified by MetaXpress® software.

### Immunohistochemistry and *in situ* hybridization of mouse embryos

Experimental procedures were performed as described previously^83, 117^. All E13.5 and E15.5 mouse embryos were fixed with 4% PFA for 2 h (for immunostaining) or 4 h (for *in situ* hybridization) and washed with 1x PBS at 4 °C. To prepare samples for 20-μm cryosectioning, the samples were cryoprotected with 30% sucrose and embedded in OCT compound (Leica). For vibratome sectioning, whole embryos were embedded in 4% low melting agarose and transversely sectioned at 300 μm. During the immunohistochemistry step, sections were permeabilized and blocked with 10% FBS plus 0.3%-0.5% Triton-X-100 for 1 h at room temperature or overnight at 4 °C. Antibodies were applied at respective titers and incubated at 4 °C for one or two nights for the 20-μm and 300-μm sections, respectively. Sections were washed frequently with wash buffer. For the 20-μm sections, samples were washed three times with 0.01% Triton X-100 in 1x PBS. For the 300-μm sections, an overnight wash with 0.5% Triton X-100 in 1x PBS (PBST) was performed. Secondary antibody was applied for 1 h at room temperature or overnight at 4 °C for the 20-μm and 300-μm sections, respectively. Finally, sections were washed with wash buffer and mounted. Primary antibodies used were: rabbit anti-Lhx3 (1:2000, Abcam Cat# ab14555, RRID:<u>AB_301332</u>); rabbit anti-Foxp1(1:20000, Abcam Cat# ab16645, RRID:<u>AB_732428</u>); goat anti-Foxp1(1:100 R&D systems Cat# AF4534, RRID:<u>AB_2107102</u>); rabbit anti-Hoxc8 (1:5000 Sigma-Aldrich Cat# HPA028911, RRID:<u>AB_10602236</u>); goat anti-Hoxd10 (1:1000 Santa-Cruz Cat# sc-33005, RRID:<u>AB_648462</u>); sheep anti-GFP (1:1000 AbD Serotec/Bio-Rad Cat# 4745–1051, RRID:<u>AB_619712</u>); rabbit anti-RFP (1:500 Abcam Cat# ab62341, RRID: <u>AB_945213</u>); mouse anti-MNR2/MNX1/HB9 (1:50 DSHB Cat# 81.5C10, RRID:<u>AB_2145209</u>); goat anti-Chat (1:100 Millipore/Sigma Cat# ab144P, RRID:<u>AB_2079751</u>); mouse anti-COUP-TF2/NR2F2 (1:200 R&D systems Cat# PP-H7147-00, RRID:<u>AB_2155627</u>); rabbit anti-Evi1/Mecom (1:500 Cell Signaling Technology Cat# 2593, RRID:<u>AB_2184098</u>); goat anti-Isl1 (1:1000 Neuromics Cat# GT15051, RRID:<u>AB_2126323</u>); mouse anti-SMI-32 (1:1000 BioLegend Cat# 801701, RRID:<u>AB_2564642</u>); rabbit anti-Satb2 (1:1000 Abcam Cat# ab92446, RRID:<u>AB_10563678</u>); guinea pig anti-Satb2 (1:1000 Synaptic Systems Cat# 327004, RRID:<u>AB_2620070</u>); rat anti-Bcl11b/Ctip2 (1:1000 Abcam Cat# ab18465, RRID:<u>AB_2064130</u>); rabbit anti-Zfhx4 (1:200 Novus Cat# NBP1-82156, RRID:<u>AB_11020060</u>); rabbit anti-nNos (1:10000 Immunostar Cat# 24287, RRID:<u>AB_572256</u>); sheep anti-Onecut2 (1:500 Novus Cat# AF6294, RRID:<u>AB_10640365</u>); rabbit anti-Calbindin D-28K/Calb1 (1:1000 Millipore/Sigma Cat# AB1778, RRID:<u>AB_2068336</u>); rabbit anti-Nfib (1:1000 Novus Cat# NBP1-81000, RRID:<u>AB_11027763</u>); rabbit anti-Grm5 (1:500 Millipore/Sigma Cat# AB5675, RRID:<u>AB_2295173</u>); sheep anti-Ebf2 (1:500 Novus Cat# AF7006, RRID:<u>AB_10972102</u>); rabbit anti-cleaved caspase-3 (1:500 Cell Signaling Cat# 9661, RRID:<u>AB_2341188</u>). Guinea pig anti-Foxp1 (1:320000), anti-Pou3f1 (1:2000, cat# CU822, RRID:<u>AB_2631303</u>), anti-Isl1 (1:10000, cat# CU1277, RRID:<u>AB_2631974</u>), anti-Lhx1 (1:20000, cat# CU453, RRID:<u>AB_2827967</u>) were antibody gifts from Thomas Jessell and guinea pig anti-Mnx1 (1:1000) was gift from Hynek Wichterle. Guinea pig anti-Hoxa5 (1:20000, RRID:<u>AB_2744661</u>) was made in house. Alexa 488-, Cy3- and Cy5-conjugated secondary antibodies (Jackson ImmunoResearch Lab or Invitrogen Antibodies) were used at dilution titer of 1:1000.

For *in situ* hybridization, sections were dried, post-fixed with 4% PFA for 15 min and rinsed with 1x PBS at room temperature. 3 μg/ml Proteinase K treatment was applied for 5 min. Slides were then rinsed twice with 1x PBS and then acetylated for 10 min before washing.

Prehybridization was performed for at least 2 h. Riboprobes (150 ng) were heat-denatured at 80 °C for 5 min and hybridized to sections overnight at 58 °C. After washing and blocking, the slides were incubated with anti-digoxigenin-AP, Fab fragments overnight at 4 °C. After washing, the slides were color-developed with NBT/BCIP solution. Sequences for riboprobe generation are indicated in Supplementary Table 11, and the template DNA were amplified either by cloning or polymerase chain reaction.

All images were acquired using either Zeiss LSM780 or LSM980 confocal microscopes and an AxioImager Z1 upright fluorescence microscope. Images of axonal tracing are projections of z-stacks.

### RNAScope

For *Sst* and *Npy* colocalization analysis, transcripts were detected using RNAscope Multiplex Fluorescent Reagent Kit v2 (Cat. No. 323100) and probes were detected with Akoya Biosciences Opal 520 (Cat. No. FP1487001KT) and 570 (FP1488001KT). The staining protocol followed the manufacturer’s recommendations with minor modification. E13.5 embryos were fixed at 4% PFA for 30 h at 4 °C and washed with 1x PBS at 4 °C. Embedded samples were subjected to 14-μm cryosectioning. The fresh-frozen protocol was adapted by skipping the 15 min postfix stage and instead proceeding directly with pretreatment and the RNAscope hybridization assay. Adjacent slides were immunostained for spatial genes to determine the position of signals (rostral: *Hoxa5*; caudal: *Hoxc8*; medial: *Foxp1^+^ Isl1^+^*; lateral: *Foxp1^+^ Isl1^-^*). More details of probes against *Sst* (ACD Cat# 404631) and *Npy* (ACD Cat# 313321-C2) are presented in Supplementary Table 11.

### Human embryonic spinal cord sample collection and immunostaining

Human embryonic spinal cords were collected as described previously^79^. Fetal embryos at gestation week 6.3 and 10.3 were harvested from the Department of Gynecology and Obstetrics at Antoine Béclère Hospital (Clamart, France), with appropriate maternal written consent for scientific use and approval from the local Medical Ethics Committee and by the French Biomedicine Agency (reference number PFS 12-002). No pregnancy had been terminated due to fetal abnormality and all pregnant mothers had undergone legally-induced abortions during the first trimester of pregnancy. Fetal limb and feet lengths were measured for fetal age determination^118^.

All spinal cords were fixed with fresh cold 4% PFA for 90 min, before being washed and equilibrated overnight with 30% sucrose. Samples were then embedded in OCT and subjected to 16-μm cryosectioning. For immunostaining, spinal cord sections were blocked and permeabilized with 10% FBS and 0.2% Triton X-100 prepared in 1x PBS for 10 min at room temperature. Sections were incubated with primary antibodies diluted to an appropriate titer in 2% FBS and 0.2% Triton X-100 prepared in 1x PBS. Sections were washed three times in 1x PBS before secondary antibodies were applied for 1 h at room temperature. Finally, sections were washed and mounted in Fluoromount (Sigma-Aldrich or Cliniscience). All images were acquired with a DM6000 microscope (Leica) and a CoolSNAP EZ CDD camera.

### Sample collection and *in situ* hybridization of zebrafish embryos

Adult zebrafish, *Danio rerio*, were obtained from Dr. Sheng-Ping Hwang at ICOB, Academia Sinica. Embryos were collected and maintained as described previously^119^. From 22 h post-fertilization (hpf), the embryos were raised in 0.2 mM PTU (1-phenyl-2-thiourea) at 28 °C to block pigmentation and to permit visualization, before being fixed overnight with 4% PFA in PBST (0.1% Tween-20 in 1xPBS) at 4 °C. The fixed embryos were washed with 1x PBST, dehydrated with methanol, and then stored at −20 °C until subjected to *in situ* hybridization (ISH). The PCR primers used to amplify cDNAs for probe synthesis were designed based on Zfin database sequences^120^ and are listed in Supplementary Table 11. ISH was performed following standard procedures^121^. For dual-fluorescence *in situ* hybridization (FISH), the embryos were hybridized with DIG-labeled RNA probe (Roche) and DNP-labeled RNA probe (Mirus Bio), and signals were detected using a Tyramide Signal Amplification (TSA) Plus kit (PerkinElmer). The embryos were imaged using a Zeiss Axio Imager A2 microscope or a Zeiss LSM 880 with an Airyscan confocal system.

### Analysis of the spatial distribution of MMC MN subpopulations

The positioning of MMC MN subpopulations in E13.5 brachial spinal segments was analyzed similarly to previous descriptions^103, 122^. Coordinates (x,y) were assigned based on the position of each MMC MN using the “spots” function in the imaging software Imaris 9.5.1 (Bitplane). Four additional coordinates were exported, i.e. midpoints for the dorsoventral and mediolateral spinal cord boundary. To account for differences in spinal cord size and shape, sections were normalized to a standardized hemi-section spinal cord (midline to lateral = 400 μm; dorsal to ventral edge = 800 μm). Density distributions for cells were plotted using the ggplot2 ‘geom_density’ function.

## Statistical analysis

Statistical analysis was performed in GraphPad Prism 6.0 (GraphPad Software). One-way ANOVAs with Tukey’s multiple comparison test was used to analyze multiple experimental groups. Two groups were analyzed by Student’s or paired *t*-tests.Values are presented as mean ± SD (standard deviation). Statistical significance is represented as * p<0.05 or ** p<0.01, unless indicated otherwise.

## Data and code availability

The E13.5 mouse spinal MNs scRNA-seq raw data generated for this study is available in the GEO database, which will be released for download upon publication. The codes used for scRNA-seq data analysis were an adaptation of standard R packages, as described in the Methods section. The codes used for estimating the number of clusters and performing similarity analyses between cell groups are available on GitHub at https://github.com/sqjin/MotorNeuron. More detailed information is available upon request.

## Acknowledgements

Nr2f2 floxed mice were generously provided by Ming-Jer Tsai (Baylor College of Medicine) and Li-Ru You (National Yang Ming Chiao Tung University). We also thank Hynek Wichterle, Tom Jessell, and Susan Morton (Columbia University) for many gifts of antibodies used in this study. We acknowledge the FACS, Transgenics, Genomics, Bioinformatic and Imaging cores of IMB Academia Sinica for their strong technical assistance, and the IMB Scientific English Editing Core for further editing the manuscript. We thank Shen-Ju Chou and Suewei Lin (Academia Sinica) for valuable inputs and members of the JAC lab for discussion and proofreading. This work was supported by Academia Sinica (CDA-107-L05 and AS-GC-109–03), MOST (110-2326-B-001-009, 109-2314-B-001-010-MY3, 108–2311-B-001–011) and NHRI (NHRI-EX110-10831NI) (J.-A.C.); a NSF grant DMS1763272 and a Simons Foundation grant (594598, Q.N.).

## Author contributions

Conceptualization: J.-A.C. and E.S.L.; Computational analysis: S.J., E.S.L., Y.-C.C.; Experimental work: E.S.L., W.-S.L., L.W.Y., C.-T.T., M.C.; Writing – Original Draft: J.-A.C., E.S.L.; Writing – Review & Editing: J.-A.C., E.S.L., S.J., Y.-C.C., S.N., J.-K.Y., Y.-H.S., Q.N.; Funding Acquisition: J.-A.C.; Supervision: J.-A.C. and Q.N.

## Supplemental information

Supplementary Table 1: Marker genes for all rostral MN clusters (related to Fig. 1e).

Supplementary Table 2: Marker genes for all caudal MN clusters (related to Fig. 1f).

Supplementary Table 3: Differentially expressed genes between rostral LMC and MMC MNs (related to Extended Data Fig. 4b).

Supplementary Table 4: Differentially expressed genes between caudal LMC and MMC MNs (related to Extended Data Fig. 4b).

Supplementary Table 5: Differentially expressed genes between rostral and caudal brachial LMC subclusters (related to Fig. 3d).

Supplementary Table 6: Differentially expressed genes between medial and lateral LMC subclusters in brachial segments (related to Fig. 3e).

Supplementary Table 7: Marker genes for each brachial LMC subcluster (related to Fig. 3 and Extended Data Fig. 5).

Supplementary Table 8: Highly variable genes across brachial LMC MNs.

Supplementary Table 9: Marker genes for each lumbar LMC subcluster (related to Extended Data Fig. 7).

Supplementary Table 10: Differentially expressed genes between brachial and lumbar LMC MNs (related to Extended Data Fig. 8).

Supplementary Table 11: *In situ* hybridization probe primers used in this study (Related to Fig. 2b, 4c, 5, 6g, 7 and Extended Data Fig. 4c).

Supplementary Table 12: qPCR primer used in this study (Related to Extended Data Fig. 1b, c).

## References

1. Osseward, P. J. & Pfaff, S. L. Cell type and circuit modules in the spinal cord. Curr Opin Neurobiol 56, 175–184, doi:10.1016/j.conb.2019.03.003 (2019).

2. Delile, J. et al. Single cell transcriptomics reveals spatial and temporal dynamics of gene expression in the developing mouse spinal cord. Development 146, doi:10.1242/dev.173807 (2019).

3. Stifani, N. Motor neurons and the generation of spinal motor neuron diversity. Front Cell Neurosci 8, 293, doi:10.3389/fncel.2014.00293 (2014).

4. Sagner, A. & Briscoe, J. Establishing neuronal diversity in the spinal cord: a time and a place. Development 146, doi:10.1242/dev.182154 (2019).

5. O’Reilly, J. C., Summers, A. P. & Ritter, D. A. The Evolution of the Functional Role of Trunk Muscles During Locomotion in Adult Amphibians1. American Zoologist 40, 123–135, doi:10.1093/icb/40.1.123 (2000).

6. D’Elia, K. P. & Dasen, J. S. Development, functional organization, and evolution of vertebrate axial motor circuits. Neural Dev 13, 10, doi:10.1186/s13064-018-0108-7 (2018).

7. Fetcho, J. R. A review of the organization and evolution of motoneurons innervating the axial musculature of vertebrates. Brain Res 434, 243–280 (1987).

8. De Marco Garcia, N. V. & Jessell, T. M. Early motor neuron pool identity and muscle nerve trajectory defined by postmitotic restrictions in Nkx6.1 activity. Neuron 57, 217–231, doi:10.1016/j.neuron.2007.11.033 (2008).

9. Livet, J. et al. ETS gene Pea3 controls the central position and terminal arborization of specific motor neuron pools. Neuron 35, 877–892, doi:10.1016/s0896-6273(02)00863-2 (2002).

10. Dasen, J. S., Tice, B. C., Brenner-Morton, S. & Jessell, T. M. A Hox regulatory network establishes motor neuron pool identity and target-muscle connectivity. Cell 123, 477–491 (2005).

11. Shirasaki, R., Lewcock, J. W., Lettieri, K. & Pfaff, S. L. FGF as a target-derived chemoattractant for developing motor axons genetically programmed by the LIM code. Neuron 50, 841–853, doi:10.1016/j.neuron.2006.04.030 (2006).

12. Pecho-Vrieseling, E., Sigrist, M., Yoshida, Y., Jessell, T. M. & Arber, S. Specificity of sensory-motor connections encoded by Sema3e-Plxnd1 recognition. Nature 459, 842–846, doi:10.1038/nature08000 (2009).

13. Vrieseling, E. & Arber, S. Target-induced transcriptional control of dendritic patterning and connectivity in motor neurons by the ETS gene Pea3. Cell 127, 1439–1452, doi:10.1016/j.cell.2006.10.042 (2006).

14. Helmbacher, F., Schneider-Maunoury, S., Topilko, P., Tiret, L. & Charnay, P. Targeting of the EphA4 tyrosine kinase receptor affects dorsal/ventral pathfinding of limb motor axons. Development 127, 3313–3324 (2000).

15. Huber, A. B. et al. Distinct roles for secreted semaphorin signaling in spinal motor axon guidance. Neuron 48, 949–964, doi:10.1016/j.neuron.2005.12.003 (2005).

16. Tang, F. et al. mRNA-Seq whole-transcriptome analysis of a single cell. Nat Methods 6, 377–382, doi:10.1038/nmeth.1315 (2009).

17. Macosko, E. Z. et al. Highly Parallel Genome-wide Expression Profiling of Individual Cells Using Nanoliter Droplets. Cell 161, 1202–1214, doi:10.1016/j.cell.2015.05.002 (2015).

18. Consortium, T. M. et al. Single-cell transcriptomics of 20 mouse organs creates a Tabula Muris. Nature 562, 367–372, doi:10.1038/s41586-018-0590-4 (2018).

19. Han, X. et al. Construction of a human cell landscape at single-cell level. Nature 581, 303–309, doi:10.1038/s41586-020-2157-4 (2020).

20. Briscoe, J. & Marín, O. Looking at neurodevelopment through a big data lens. Science 369, doi:10.1126/science.aaz8627 (2020).

21. Ozel, M. N. et al. Neuronal diversity and convergence in a visual system developmental atlas. Nature 589, 88–95, doi:10.1038/s41586-020-2879-3 (2021).

22. Maniatis, S. et al. Spatiotemporal dynamics of molecular pathology in amyotrophic lateral sclerosis. Science 364, 89–93, doi:10.1126/science.aav9776 (2019).

23. Ren, X. et al. COVID-19 immune features revealed by a large-scale single-cell transcriptome atlas. Cell 184, 1895–1913.e1819, doi:10.1016/j.cell.2021.01.053 (2021).

24. Bergen, V., Lange, M., Peidli, S., Wolf, F. A. & Theis, F. J. Generalizing RNA velocity to transient cell states through dynamical modeling. Nat Biotechnol 38, 1408–1414, doi:10.1038/s41587-020-0591-3 (2020).

25. Di Bella, D. J. et al. Molecular logic of cellular diversification in the mouse cerebral cortex. Nature 595, 554–559, doi:10.1038/s41586-021-03670-5 (2021).

26. Blum, J. A. et al. Single-cell transcriptomic analysis of the adult mouse spinal cord reveals molecular diversity of autonomic and skeletal motor neurons. Nat Neurosci 24, 572–583, doi:10.1038/s41593-020-00795-0 (2021).

27. Alkaslasi, M. R. et al. Single nucleus RNA-sequencing defines unexpected diversity of cholinergic neuron types in the adult mouse spinal cord. Nat Commun 12, 2471, doi:10.1038/s41467-021-22691-2 (2021).

28. Ladle, D. R., Pecho-Vrieseling, E. & Arber, S. Assembly of motor circuits in the spinal cord: driven to function by genetic and experience-dependent mechanisms. Neuron 56, 270–283, doi:10.1016/j.neuron.2007.09.026 (2007).

29. Briggs, J. A. et al. Mouse embryonic stem cells can differentiate via multiple paths to the same state. Elife 6, doi:10.7554/eLife.26945 (2017).

30. Satija, R., Farrell, J. A., Gennert, D., Schier, A. F. & Regev, A. Spatial reconstruction of single-cell gene expression data. Nat Biotechnol 33, 495–502, doi:10.1038/nbt.3192 (2015).

31. Hanley, O. et al. Parallel Pbx-Dependent Pathways Govern the Coalescence and Fate of Motor Columns. Neuron 91, 1005–1020, doi:10.1016/j.neuron.2016.07.043 (2016).

32. Chen, T. H. & Chen, J. A. Multifaceted roles of microRNAs: From motor neuron generation in embryos to degeneration in spinal muscular atrophy. Elife 8, doi:10.7554/eLife.50848 (2019).

33. Velasco, S. et al. A Multi-step Transcriptional and Chromatin State Cascade Underlies Motor Neuron Programming from Embryonic Stem Cells. Cell Stem Cell 20, 205–217.e208, doi:10.1016/j.stem.2016.11.006 (2017).

34. Hinckley, C. A., Hartley, R., Wu, L., Todd, A. & Ziskind-Conhaim, L. Locomotor-like rhythms in a genetically distinct cluster of interneurons in the mammalian spinal cord. J Neurophysiol 93, 1439–1449, doi:10.1152/jn.00647.2004 (2005).

35. Stam, F. J. et al. Renshaw cell interneuron specialization is controlled by a temporally restricted transcription factor program. Development 139, 179–190, doi:10.1242/dev.071134 (2012).

36. Dasen, J. S., De Camilli, A., Wang, B., Tucker, P. W. & Jessell, T. M. Hox repertoires for motor neuron diversity and connectivity gated by a single accessory factor, FoxP1. Cell 134, 304-316, doi:10.1016/j.cell.2008.06.019 (2008).

37. Huang, T. et al. Identifying the pathways required for coping behaviours associated with sustained pain. Nature 565, 86–90, doi:10.1038/s41586-018-0793-8 (2019).

38. Gutierrez-Mecinas, M. et al. Preprotachykinin A is expressed by a distinct population of excitatory neurons in the mouse superficial spinal dorsal horn including cells that respond to noxious and pruritic stimuli. Pain 158, 440–456, doi:10.1097/j.pain.0000000000000778 (2017).

39. Gore, B. B., Wong, K. G. & Tessier-Lavigne, M. Stem cell factor functions as an outgrowth-promoting factor to enable axon exit from the midline intermediate target. Neuron 57, 501–510, doi:10.1016/j.neuron.2008.01.006 (2008).

40. Hirata, T. et al. Stem cell factor induces outgrowth of c-kit-positive neurites and supports the survival of c-kit-positive neurons in dorsal root ganglia of mouse embryos. Development 119, 49–56 (1993).

41. Schaller, S. et al. Novel combinatorial screening identifies neurotrophic factors for selective classes of motor neurons. Proc Natl Acad Sci U S A 114, E2486–E2493, doi:10.1073/pnas.1615372114 (2017).

42. Pischedda, F. et al. A cell surface biotinylation assay to reveal membrane-associated neuronal cues: Negr1 regulates dendritic arborization. Mol Cell Proteomics 13, 733–748, doi:10.1074/mcp.M113.031716 (2014).

43. Hashimoto, T., Maekawa, S. & Miyata, S. IgLON cell adhesion molecules regulate synaptogenesis in hippocampal neurons. Cell Biochem Funct 27, 496–498, doi:10.1002/cbf.1600 (2009).

44. Gil, O. D., Zanazzi, G., Struyk, A. F. & Salzer, J. L. Neurotrimin mediates bifunctional effects on neurite outgrowth via homophilic and heterophilic interactions. J Neurosci 18, 9312–9325 (1998).

45. Tsuchida, T. et al. Topographic organization of embryonic motor neurons defined by expression of LIM homeobox genes. Cell 79, 957–970 (1994).

46. Catela, C., Shin, M. M., Lee, D. H., Liu, J. P. & Dasen, J. S. Hox Proteins Coordinate Motor Neuron Differentiation and Connectivity Programs through Ret/Gfralpha Genes. Cell Rep 14, 1901–1915, doi:10.1016/j.celrep.2016.01.067 (2016).

47. Poliak, S. et al. Synergistic integration of Netrin and ephrin axon guidance signals by spinal motor neurons. Elife 4, doi:10.7554/eLife.10841 (2015).

48. Bonanomi, D. & Pfaff, S. L. Motor axon pathfinding. Cold Spring Harb Perspect Biol 2, a001735, doi:10.1101/cshperspect.a001735 (2010).

49. Catela, C. & Kratsios, P. Transcriptional mechanisms of motor neuron development in vertebrates and invertebrates. Dev Biol 475, 193–204, doi:10.1016/j.ydbio.2019.08.022 (2021).

50. Price, S. R., De Marco Garcia, N. V., Ranscht, B. & Jessell, T. M. Regulation of motor neuron pool sorting by differential expression of type II cadherins. Cell 109, 205–216, doi:10.1016/s0092-8674(02)00695-5 (2002).

51. Lacombe, J. et al. Genetic and functional modularity of Hox activities in the specification of limb-innervating motor neurons. PLoS Genet 9, e1003184, doi:10.1371/journal.pgen.1003184 (2013).

52. Lamballe, F. et al. Pool-specific regulation of motor neuron survival by neurotrophic support. J Neurosci 31, 11144–11158, doi:10.1523/JNEUROSCI.2198-11.2011 (2011).

53. Helmbacher, F. et al. Met signaling is required for recruitment of motor neurons to PEA3-positive motor pools. Neuron 39, 767–777, doi:10.1016/s0896-6273(03)00493-8 (2003).

54. Kanning, K. C., Kaplan, A. & Henderson, C. E. Motor neuron diversity in development and disease. Annu Rev Neurosci 33, 409–440, doi:10.1146/annurev.neuro.051508.135722 (2010).

55. Hirono, K., Kohwi, M., Clark, M. Q., Heckscher, E. S. & Doe, C. Q. The Hunchback temporal transcription factor establishes, but is not required to maintain, early-born neuronal identity. Neural Dev 12, 1, doi:10.1186/s13064-017-0078-1 (2017).

56. Nusbaum, M. P., Blitz, D. M. & Marder, E. Functional consequences of neuropeptide and small-molecule co-transmission. Nat Rev Neurosci 18, 389–403, doi:10.1038/nrn.2017.56 (2017).

57. Zeisel, A. et al. Molecular Architecture of the Mouse Nervous System. Cell 174, 999–1014.e1022, doi:10.1016/j.cell.2018.06.021 (2018).

58. Häring, M. et al. Neuronal atlas of the dorsal horn defines its architecture and links sensory input to transcriptional cell types. Nat Neurosci 21, 869–880, doi:10.1038/s41593-018-0141-1 (2018).

59. Korsunsky, I. et al. Fast, sensitive and accurate integration of single-cell data with Harmony. Nat Methods 16, 1289–1296, doi:10.1038/s41592-019-0619-0 (2019).

60. Hayashi, M. et al. Graded Arrays of Spinal and Supraspinal V2a Interneuron Subtypes Underlie Forelimb and Hindlimb Motor Control. Neuron 97, 869–884.e865, doi:10.1016/j.neuron.2018.01.023 (2018).

61. Philippidou, P. & Dasen, J. S. Hox genes: choreographers in neural development, architects of circuit organization. Neuron 80, 12–34, doi:10.1016/j.neuron.2013.09.020 (2013).

62. Hollyday, M. & Hamburger, V. An autoradiographic study of the formation of the lateral motor column in the chick embryo. Brain Res 132, 197–208, doi:10.1016/0006-8993(77)90416-4 (1977).

63. Sun, Y. G. & Chen, Z. F. A gastrin-releasing peptide receptor mediates the itch sensation in the spinal cord. Nature 448, 700–703, doi:10.1038/nature06029 (2007).

64. Liguz-Lecznar, M., Urban-Ciecko, J. & Kossut, M. Somatostatin and Somatostatin-Containing Neurons in Shaping Neuronal Activity and Plasticity. Front Neural Circuits 10, 48, doi:10.3389/fncir.2016.00048 (2016).

65. Fetcho, J. R. The spinal motor system in early vertebrates and some of its evolutionary changes. Brain Behav Evol 40, 82–97, doi:10.1159/000113905 (1992).

66. Catela, C. et al. An ancient role for collier/Olf/Ebf (COE)-type transcription factors in axial motor neuron development. Neural Dev 14, 2, doi:10.1186/s13064-018-0125-6 (2019).

67. Moret, F., Renaudot, C., Bozon, M. & Castellani, V. Semaphorin and neuropilin co-expression in motoneurons sets axon sensitivity to environmental semaphorin sources during motor axon pathfinding. Development 134, 4491–4501, doi:10.1242/dev.011452 (2007).

68. Tang, K., Rubenstein, J. L., Tsai, S. Y. & Tsai, M. J. COUP-TFII controls amygdala patterning by regulating neuropilin expression. Development 139, 1630–1639, doi:10.1242/dev.075564 (2012).

69. Alcamo, E. A. et al. Satb2 regulates callosal projection neuron identity in the developing cerebral cortex. Neuron 57, 364–377, doi:10.1016/j.neuron.2007.12.012 (2008).

70. Holland, L. Z. & Holland, N. D. Cephalochordates: A window into vertebrate origins. Curr Top Dev Biol 141, 119–147, doi:10.1016/bs.ctdb.2020.07.001 (2021).

71. Ren, Q. et al. Step-wise evolution of neural patterning by Hedgehog signalling in chordates. Nat Ecol Evol 4, 1247–1255, doi:10.1038/s41559-020-1248-9 (2020).

72. Leung, B. & Shimeld, S. M. Evolution of vertebrate spinal cord patterning. Dev Dyn 248, 1028–1043, doi:10.1002/dvdy.77 (2019).

73. Ferrier, D. E., Brooke, N. M., Panopoulou, G. & Holland, P. W. The Mnx homeobox gene class defined by HB9, MNR2 and amphioxus AmphiMnx. Dev Genes Evol 211, 103-107, doi:10.1007/s004270000124 (2001).

74. Jackman, W. R., Langeland, J. A. & Kimmel, C. B. islet reveals segmentation in the Amphioxus hindbrain homolog. Dev Biol 220, 16–26, doi:10.1006/dbio.2000.9630 (2000).

75. Pergner, J., Vavrova, A., Kozmikova, I. & Kozmik, Z. Molecular Fingerprint of Amphioxus Frontal Eye Illuminates the Evolution of Homologous Cell Types in the Chordate Retina. Front Cell Dev Biol 8, 705, doi:10.3389/fcell.2020.00705 (2020).

76. Wang, Y., Zhang, P. J., Yasui, K. & Saiga, H. Expression of Bblhx3, a LIM-homeobox gene, in the development of amphioxus Branchiostoma belcheri tsingtauense. Mech Dev 117, 315–319, doi:10.1016/s0925-4773(02)00197-1 (2002).

77. Takatori, N. & Saiga, H. Evolution of CUT class homeobox genes: insights from the genome of the amphioxus, Branchiostoma floridae. Int J Dev Biol 52, 969–977, doi:10.1387/ijdb.072541nt (2008).

78. Seredick, S., Hutchinson, S. A., Van Ryswyk, L., Talbot, J. C. & Eisen, J. S. Lhx3 and Lhx4 suppress Kolmer-Agduhr interneuron characteristics within zebrafish axial motoneurons. Development 141, 3900–3909, doi:10.1242/dev.105718 (2014).

79. Mouilleau, V. et al. Dynamic extrinsic pacing of the HOX clock in human axial progenitors controls motor neuron subtype specification. Development 148, doi:10.1242/dev.194514 (2021).

80. Rayon, T., Maizels, R. J., Barrington, C. & Briscoe, J. Single cell transcriptome profiling of the human developing spinal cord reveals a conserved genetic programme with human specific features. Development, doi:10.1242/dev.199711 (2021).

81. Osseward, P. J. et al. Conserved genetic signatures parcellate cardinal spinal neuron classes into local and projection subsets. Science 372, 385–393, doi:10.1126/science.abe0690 (2021).

82. Chen, J. A. & Wichterle, H. Apoptosis of limb innervating motor neurons and erosion of motor pool identity upon lineage specific dicer inactivation. Front Neurosci 6, 69, doi:10.3389/fnins.2012.00069 (2012).

83. Tung, Y. T. et al. Mir-17∼92 Confers Motor Neuron Subtype Differential Resistance to ALS-Associated Degeneration. Cell Stem Cell 25, 193–209.e197, doi:10.1016/j.stem.2019.04.016 (2019).

84. Li, C. J. et al. MicroRNA filters Hox temporal transcription noise to confer boundary formation in the spinal cord. Nat Commun 8, 14685, doi:10.1038/ncomms14685 (2017).

85. Li, C. J. et al. MicroRNA governs bistable cell differentiation and lineage segregation via a noncanonical feedback. Mol Syst Biol 17, e9945, doi:10.15252/msb.20209945 (2021).

86. Carr, P. A., Alvarez, F. J., Leman, E. A. & Fyffe, R. E. Calbindin D28k expression in immunohistochemically identified Renshaw cells. Neuroreport 9, 2657–2661, doi:10.1097/00001756-199808030-00043 (1998).

87. Fahandejsaadi, A., Leung, E., Rahaii, R., Bu, J. & Geula, C. Calbindin-D28K, parvalbumin and calretinin in primate lower motor neurons. Neuroreport 15, 443–448, doi:10.1097/00001756-200403010-00012 (2004).

88. Spruill, M. M. & Kuncl, R. W. Calbindin-D28K is increased in the ventral horn of spinal cord by neuroprotective factors for motor neurons. J Neurosci Res 93, 1184–1191, doi:10.1002/jnr.23562 (2015).

89. Copani, A. et al. The metabotropic glutamate receptor mGlu5 controls the onset of developmental apoptosis in cultured cerebellar neurons. Eur J Neurosci 10, 2173–2184, doi:10.1046/j.1460-9568.1998.00230.x (1998).

90. Tomiyama, M. et al. Expression of metabotropic glutamate receptor mRNAs in the human spinal cord: implications for selective vulnerability of spinal motor neurons in amyotrophic lateral sclerosis. J Neurol Sci 189, 65–69, doi:10.1016/s0022-510x(01)00561-5 (2001).

91. Schellino, R., Boido, M. & Vercelli, A. The Dual Nature of Onuf’s Nucleus: Neuroanatomical Features and Peculiarities, in Health and Disease. Front Neuroanat 14, 572013, doi:10.3389/fnana.2020.572013 (2020).

92. Romanes, G. J. The motor cell columns of the lumbo-sacral spinal cord of the cat. J Comp Neurol 94, 313–363 (1951).

93. Landmesser, L. The distribution of motoneurones supplying chick hind limb muscles. J Physiol 284, 371–389 (1978).

94. Hobert, O. Regulatory logic of neuronal diversity: terminal selector genes and selector motifs. Proc Natl Acad Sci U S A 105, 20067–20071, doi:10.1073/pnas.0806070105 (2008).

95. Patel, T., Hammelman, J., Closser, M., Gifford, D. K. & Wichterle, H. General and cell-type-specific aspects of the motor neuron maturation transcriptional program. bioRxiv, 2021.2003.2005.434185, doi:10.1101/2021.03.05.434185 (2021).

96. Hobert, O. Homeobox genes and the specification of neuronal identity. Nat Rev Neurosci, doi:10.1038/s41583-021-00497-x (2021).

97. Villar, M. J. et al. Immunoreactive Calcitonin Gene-Related Peptide, Vasoactive Intestinal Polypeptide, and Somatostatin in Developing Chicken Spinal Cord Motoneurons. Eur J Neurosci 1, 269–287, doi:10.1111/j.1460-9568.1989.tb00795.x (1989).

98. New, H. V. & Mudge, A. W. Calcitonin gene-related peptide regulates muscle acetylcholine receptor synthesis. Nature 323, 809–811, doi:10.1038/323809a0 (1986).

99. Perkel, J. M. Single-cell analysis enters the multiomics age. Nature 595, 614–616, doi:10.1038/d41586-021-01994-w (2021).

100. Jin, S., Zhang, L. & Nie, Q. scAI: an unsupervised approach for the integrative analysis of parallel single-cell transcriptomic and epigenomic profiles. Genome Biol 21, 25, doi:10.1186/s13059-020-1932-8 (2020).

101. Gutman, C. R., Ajmera, M. K. & Hollyday, M. Organization of motor pools supplying axial muscles in the chicken. Brain Res 609, 129–136, doi:10.1016/0006-8993(93)90865-k (1993).

102. Kanai, M. I., Okabe, M. & Hiromi, Y. seven-up Controls switching of transcription factors that specify temporal identities of Drosophila neuroblasts. Dev Cell 8, 203–213, doi:10.1016/j.devcel.2004.12.014 (2005).

103. Chang, S. H., Su, Y. C., Chang, M. & Chen, J. A. MicroRNAs mediate precise control of spinal interneuron populations to exert delicate sensory-to-motor outputs. Elife 10, doi:10.7554/eLife.63768 (2021).

104. Jung, H. & Dasen, J. S. Evolution of patterning systems and circuit elements for locomotion. Dev Cell 32, 408–422, doi:10.1016/j.devcel.2015.01.008 (2015).

105. Dasen, J. S. & Jessell, T. M. Hox networks and the origins of motor neuron diversity. Curr Top Dev Biol 88, 169–200, doi:10.1016/S0070-2153(09)88006-X (2009).

106. Hutchinson, S. A. & Eisen, J. S. Islet1 and Islet2 have equivalent abilities to promote motoneuron formation and to specify motoneuron subtype identity. Development 133, 2137–2147, doi:10.1242/dev.02355 (2006).

107. Seredick, S. D., Van Ryswyk, L., Hutchinson, S. A. & Eisen, J. S. Zebrafish Mnx proteins specify one motoneuron subtype and suppress acquisition of interneuron characteristics. Neural Dev 7, 35, doi:10.1186/1749-8104-7-35 (2012).

108. Jung, H. et al. The Ancient Origins of Neural Substrates for Land Walking. Cell 172, 667–682.e615, doi:10.1016/j.cell.2018.01.013 (2018).

109. Wichterle, H., Lieberam, I., Porter, J. A. & Jessell, T. M. Directed differentiation of embryonic stem cells into motor neurons. Cell 110, 385–397 (2002).

110. Dessaud, E. et al. Interpretation of the sonic hedgehog morphogen gradient by a temporal adaptation mechanism. Nature 450, 717–720, doi:10.1038/nature06347 (2007).

111. Hippenmeyer, S. et al. A developmental switch in the response of DRG neurons to ETS transcription factor signaling. PLoS Biol 3, e159, doi:10.1371/journal.pbio.0030159 (2005).

112. Takamoto, N. et al. COUP-TFII is essential for radial and anteroposterior patterning of the stomach. Development 132, 2179–2189, doi:10.1242/dev.01808 (2005).

113. von Luxburg, U. A tutorial on spectral clustering. Statistics and Computing 17, 395–416, doi:10.1007/s11222-007-9033-z (2007).

114. Hu, Z., Ahmed, A. A. & Yau, C. An interpretable meta-clustering framework for single-cell RNA-Seq data integration and evaluation. bioRxiv, 2021.2003.2029.437525, doi:10.1101/2021.03.29.437525 (2021).

115. Gu, Z., Eils, R. & Schlesner, M. Complex heatmaps reveal patterns and correlations in multidimensional genomic data. Bioinformatics 32, 2847–2849, doi:10.1093/bioinformatics/btw313 (2016).

116. Yu, G., Wang, L.-G., Han, Y. & He, Q.-Y. clusterProfiler: an R Package for Comparing Biological Themes Among Gene Clusters. OMICS: A Journal of Integrative Biology 16, 284–287, doi:10.1089/omi.2011.0118 (2012).

117. Chen, J. A. et al. Mir-17-3p controls spinal neural progenitor patterning by regulating Olig2/Irx3 cross-repressive loop. Neuron 69, 721–735, doi:10.1016/j.neuron.2011.01.014 (2011).

118. Evtouchenko, L., Studer, L., Spenger, C., Dreher, E. & Seiler, R. W. A mathematical model for the estimation of human embryonic and fetal age. Cell Transplant 5, 453–464, doi:10.1016/0963-6897(96)00079-6 (1996).

119. Westerfield, M. The zebrafish book. A guide for the laboratory use of zebrafish (Danio rerio). 4th edn, (Univ. of Oregon Press, Eugene, 2000).

120. Ruzicka, L. et al. The Zebrafish Information Network: new support for non-coding genes, richer Gene Ontology annotations and the Alliance of Genome Resources. Nucleic Acids Res 47, D867–D873, doi:10.1093/nar/gky1090 (2019).

121. Thisse, C. & Thisse, B. High-resolution in situ hybridization to whole-mount zebrafish embryos. Nat Protoc 3, 59–69, doi:10.1038/nprot.2007.514 (2008).

122. Dewitz, C. et al. Nuclear Organization in the Spinal Cord Depends on Motor Neuron Lamination Orchestrated by Catenin and Afadin Function. Cell Rep 22, 1681–1694, doi:10.1016/j.celrep.2018.01.059 (2018).

